# Developmental exposure to domoic acid disrupts startle response behavior and circuitry

**DOI:** 10.1101/2021.01.08.425996

**Authors:** Jennifer M. Panlilio, Ian T. Jones, Matthew C. Salanga, Neelakanteswar Aluru, Mark E. Hahn

## Abstract

Harmful algal blooms produce potent neurotoxins that accumulate in seafood and are hazardous to human health. Developmental exposure to the harmful algal bloom toxin, domoic acid (DomA), has behavioral consequences well into adulthood, but the cellular and molecular mechanisms are largely unknown. To assess these, we exposed zebrafish embryos to DomA during the previously identified window of susceptibility (2 days post-fertilization) and used the well-known startle response circuit as a tool to identify specific neuronal components that are targeted by exposure to DomA. Exposure to DomA reduced the probability of eliciting a startle after auditory/vibrational or electrical stimuli and led to the dramatic reduction of one type of startle, short latency c-start (SLC) responses. Furthermore, DomA-exposed larvae had altered kinematics of both SLC and long latency c-start (LLC) startle responses, exhibiting shallower bend angles and slower maximal angular velocities. Using vital dye staining, immunolabelling, and live imaging of transgenic lines, we determined that while the sensory inputs were intact, the reticulospinal neurons required for SLC responses were absent in most DomA-exposed larvae. Furthermore, axon tracing revealed that DomA-treated larvae also showed significantly reduced primary motor neuron axon collaterals. Overall, these results show that developmental exposure to DomA leads to startle deficits by targeting specific subsets of neurons. These findings provide mechanistic insights into the neurodevelopmental effects of excess glutamatergic signaling caused by exposure to DomA. It further provides a model for using the startle response circuit to identify neuronal populations targeted by toxin or toxicant exposures.

**Summary statement:** We used the zebrafish startle response as a tool to identify sensory-motor deficits and the loss of specific neural populations after developmental exposure to the harmful algal bloom toxin domoic acid.

**GRAPHICAL ABSTRACT:** 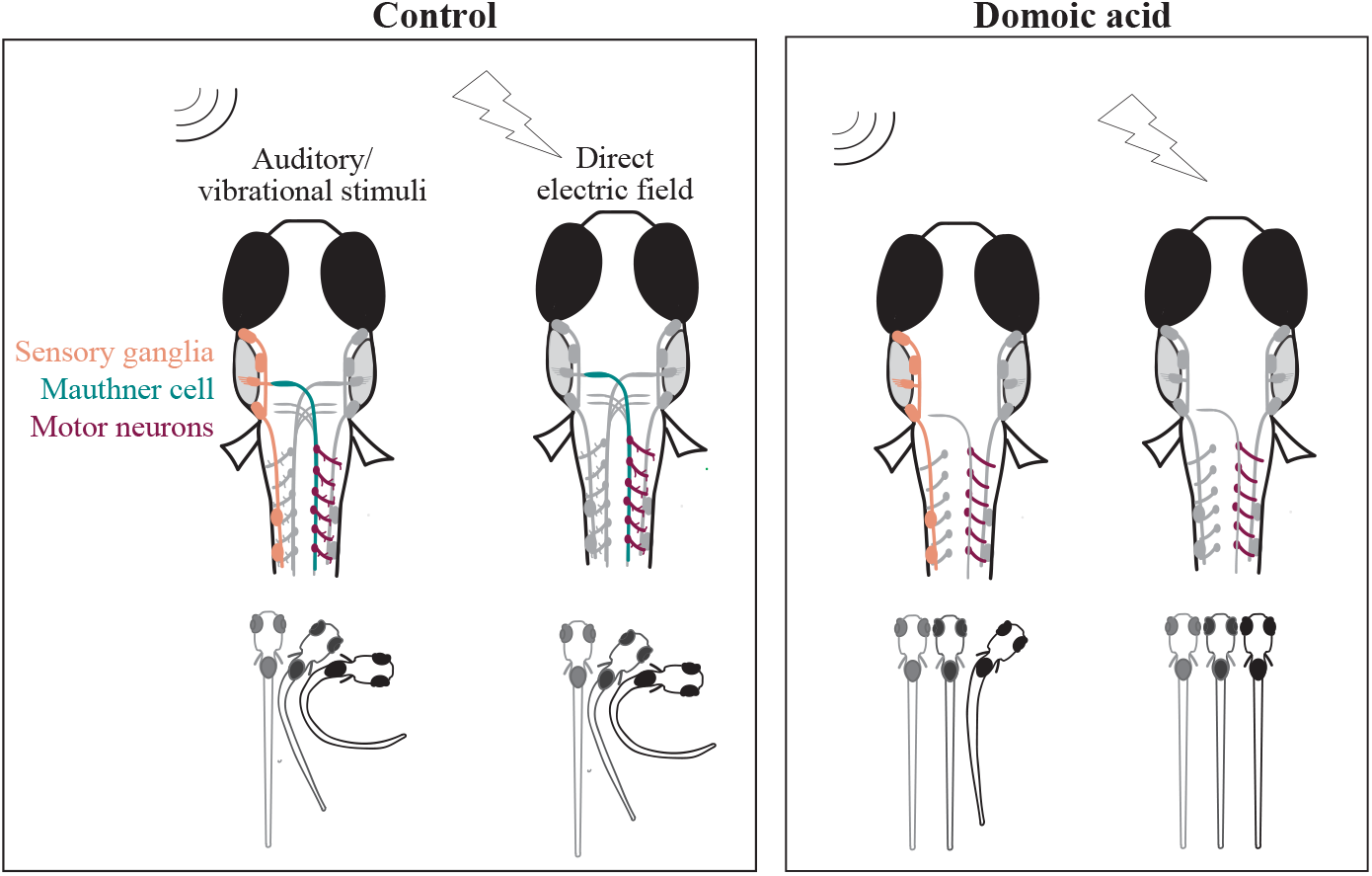

## INTRODUCTION

Harmful algal blooms (HABs) – the proliferation of algae that produce toxins – have increased in frequency, duration, and geographic extent in recent decades (Anderson et al., 2002). Some HABs produce potent toxins that contaminate drinking water, seafood, and air. One such toxin, domoic acid (DomA), is produced primarily by algae in the genus *Pseudo-nitzschia*. Increasing blooms of toxic *Pseudo-nitzschia* species, resulting in part from changing climatic conditions, have been documented in recent years on both the west and east coasts of the United States, where they have led to shellfish harvest closures (Bates et al., 2018; Clark et al., 2019; McKibben et al., 2017)

DomA is a structural analog of glutamate, and is known to exert its toxicity by binding and activating ionotrophic glutamate receptors (Cendes et al., 1995; Hampson and Manalo, 1998). Exposure to DomA in adults occurs primarily through consumption of contaminated seafood. To prevent acute toxicity to humans, a regulatory limit has been set at 20 mg of DomA per kg shellfish tissue (Wekell et al., 2010). However, this limit may not be protective for earlier life stages, which are more sensitive to DomA than adult stages (Doucette et al., 2004; Tryphonas et al., 1990; Xi et al., 1997).

DomA is an established developmental neurotoxin. It has been shown to cross the placental barrier, readily accumulate in amniotic fluids, and distribute to fetal brains in non-human mammals (Brodie et al., 2006; Lefebvre et al., 2018; Maucher and Ramsdell, 2007). This provides a means by which fetuses can continue to be exposed to DomA after it is cleared from the mother’s plasma (Brodie et al., 2006; Shum et al., 2020). Further exposures may occur in postnatal development through consumption of contaminated breast milk (Maucher and Ramsdell, 2005; Rust et al., 2014).

While regulatory limits to protect shellfish consumers from acute toxicity have been established, evidence shows that developmental exposures to doses of DomA that do not lead to acute neurotoxic phenotypes (“asymptomatic doses”) can have long-term consequences. In rodents, both prenatal and postnatal exposures to these asymptomatic doses have been shown to lead to changes in neuron receptor densities (Perry et al., 2009), altered neural connectivity (Dakshinamurti et al., 1993; Mills et al., 2016), aberrant brain morphology (Bernard et al., 2007; Doucette et al., 2004; Jian Wang et al., 2000), and ultimately behavioral deficits through adulthood. Behavioral deficits include impaired interlimb coordination (Jian Wang et al., 2000; Shiotani et al., 2017), aberrant exploratory behavior (Levin et al., 2005; Shiotani et al., 2017; Tanemura et al., 2009), reduced socialization (Mills et al., 2016; Zuloaga et al., 2016), and the inability to cope with novel environments (Doucette et al., 2004; Perry et al., 2009). Many of these behavioral and histological phenotypes are latent and progressive, either appearing for the first time or increasing in severity as the animal ages. Developmental DomA toxicity also can be silent until unmasked by further challenges such as chemical exposures in adulthood, making the animals more susceptible to developing seizures and memory deficits (Gill et al., 2010; Levin et al., 2005; Tiedeken and Ramsdell, 2007)

Although behavioral deficits resulting from low-level, developmental exposure to DomA have been extensively characterized, the cellular and molecular mechanisms that underlie these behavioral effects are poorly understood. We recently showed that exposure to DomA during a specific period in zebrafish neurodevelopment (2 dpf) can lead to myelin sheath defects and aberrant startle behavior, supporting the use of zebrafish as a model for investigating the cellular and molecular mechanisms underlying DomA-induced developmental neurotoxicity (Panlilio et al., 2020). Building on these findings by using the well-characterized startle response circuit as a tool, here we i) determine which sensory and motor processes are disrupted by DomA exposure, and ii) identify specific neuronal populations within the circuit that are disrupted by DomA toxicity.

The startle response is an appropriate tool to investigate the behavioral effects of DomA exposure. Aberrant startle response patterns have been associated with myelination defects (Pogoda et al., 2006), which have been characterized as a feature of DomA toxicity (Panlilio et al., 2020). In addition, aberrant glutamate signaling also alters startle response kinematics, further strengthening the selection of startle response as a read-out for DomA-driven glutamate excitotoxicity (McKeown et al., 2012).

Startle responses in teleosts are elicited by sudden stimuli (auditory/vibrational, tactile, visual) with different intensity levels. Auditory/vibrational stimuli can lead to either of the two types of startle responses– a short latency c (SLC)-type startle response or a long latency c (LLC)-type startle response, which are distinguished by their onset time and kinematics. SLC responses occur shortly after exposure to a stimulus (about 15 milliseconds or less, depending on temperature) and tend to lead to more pronounced bend angles, while LLCs occur later and produce shallower bend angles (Burgess and Granato, 2007a; Troconis et al., 2016). Increasing the intensity of auditory/vibrational stimuli biases fish to perform SLC responses, which requires the activation of a hindbrain neuron called the Mauthner cell (Eaton et al., 1977; Zottoli, 1977). Conversely, lower intensity stimuli are more likely to lead to LLC startles, which are Mauthner cell-independent (Jain et al., 2018; Marquart et al., 2019).

The underlying startle response circuits are well known in zebrafish, making it is possible to link startle behavioral data to underlying structural and cellular targets. Auditory/vibrational stimuli are perceived by the statoacoustic ganglia, which transmit sensory information from the hair cells in the inner ear (Faber et al., 1989; Kohashi et al., 2012). Sensory information is also perceived by the lateral line, an organ system that detects vibration and changes in flow through a series of mechanoreceptors along the body (Mirjany et al., 2011). For SLC startles, the statoacoustic ganglia cranial nerve synapses to the Mauthner cell in the hindbrain, which fires a single action potential that is propagated down its axons in the spinal cord and synapses to the primary motor neurons (Kohashi and Oda, 2008; Korn and Faber, 2005). The near-simultaneous activation of the primary motor neurons results in a ‘c’ shaped bend. In contrast to auditory/vibrational stimuli, direct-electric field stimulation bypasses the sensory system altogether, directly activating the Mauthner neurons and the downstream circuits (Tabor et al., 2014).

The goal of this study was to use this well-known startle response circuit as a tool to identify cell targets and nervous system processes that were perturbed by DomA. We show that developmental exposure to DomA reduces startle responsiveness. By varying stimulus intensities, we determine whether DomA-exposed fish have the capacity to perceive changes in sensory stimuli, as reflected by their increased responsiveness with higher stimulus intensities. By using direct electric fields, which bypass the sensory system, we show that DomA exposed fish still are unable to perform startle responses, indicating that DomA-induced startle deficits occur downstream of the sensory system. Finally, with vital dyes, transgenics, and immunolabeling, we identify specific neuronal populations in the startle circuit that are disrupted by DomA.

## METHODS

### Fish husbandry and lines used

These experiments were approved by the Woods Hole Oceanographic Institution Animal Care and Use Committee (Assurance D16-00381 from the NIH Office of Laboratory Animal Welfare). Zebrafish (*Danio rerio*) embryos were maintained at 28-28.5°C with a 14:10 light dark cycle in 0.3x Danieau’s medium. The following transgenic lines were used: *Tg*(*mbp:EGFP-CAAX*) (Almeida et al., 2011), *Tg*(*sox10:RFP*) (Kucenas et al., 2008), and *Tg*(*mbp:EGFP*) (gift from Dr. Kelly Monk, generated by Dr. Charles Kaufman in the laboratory of Dr. Leonard Zon, Harvard Medical School, Boston, MA).

### Generation of the Tg(cntn1b:EGFP-CAAX) line

The *Tg*(*cntn1b:EGFP-CAAX*) line was generated by Gibson assembly (Czopka et al., 2013). Using previously published primers, the *cntn1b* promoter region was amplified from AB wildtype fish (Czopka et al., 2013). The *cntn1b* promoter was then assembled with EGFP-CAAX (cloned out of *Tg*(*mbp:EGFP-CAAX*) fish) into the vector backbone pkHR7, which contains ISCe-1 restriction sites (Hoshijima et al., 2016). To generate the stable line, the plasmid containing *cnln1b:EGFP-CAAX* was co-injected along with I-SceI in the 1-2 cell stage (Soroldoni et al., 2009; Thermes et al., 2002). Injected fish were grown up, and their progeny that carried the transgene were used to generate embryos for experimental use.

### Domoic acid exposure paradigm (Panlilio et al., 2020)

Domoic acid (5 mg; Sigma-Aldrich, MO) was dissolved directly in the vial with diluted embryo medium (0.2x Danieau’s) to obtain a 20 mM solution which was immediately used to generate stock concentrations of 0.675 μg/μl and 1.4 μg/μl. Aliquots were then stored at −20°C. Working solutions were prepared fresh prior to microinjection by diluting the stock in 0.2x Danieau’s to obtain the appropriate doses.

Microinjection needles were prepared from glass capillary tubes (0.58 mm inner diameter; World Precision Instruments, FL; 1B100F-4) using a pipette puller (Sutter instrument model p-30, heat 750, pull= 0). Microinjections were then performed using Narishige IM-300 microinjector, which was calibrated to consistently deliver 0.2-nL by adjusting gate time (milliseconds) and back pressure.

DomA (nominal concentration of 0.14 ng) was intravenously microinjected into the common posterior cardinal vein at 48-52 hpf (Panlilio et al., 2020). Controls from the same breeding clutch were injected with the saline vehicle (0.2x Danieau’s). To perform intravenous microinjections, fish were anesthetized with 0.10% w/v Tricaine mesylate (MS222), and placed laterally on dishes coated with 1.5% agarose (Cianciolo Cosentino et al., 2010). An injection was deemed successful if there was a visible displacement of blood cells and no disruption of the yolk during the injection process. Any fish that showed evidence of being incorrectly injected were immediately removed from the study. Following injections, zebrafish were placed back in clean embryo media and monitored daily.

## Measuring Startle Responses

Two different types of stimuli were used to elicit startle behavior: auditory/vibrational (A/V) and direct electric field stimuli.

### Auditory/ Vibrational (A/V) stimulation

A/V stimuli were generated by a minishaker (Brüel & Kjaer, Vibration Exciter 4810) connected to an amplifier (Brüel & Kjaer, Power Amplifier Type 2718) (Burgess and Granato, 2007b; Wolman et al., 2011). Larvae were tested in a 16-well acrylic plate (40 x 40 mm) that rested in a petri dish (100mm x 100mm). The 16-well plate itself comprised of laser cut acrylic pieces that were fused together using acrylic cement (Weld-On #3; IPS) (Wolman et al., 2011). An LED backlight was placed below the dish to illuminate the well plate (Adafruit #1622). The petri dish was epoxied to a thumbscrew (#10-32 UNF threads) that was fitted to the base of the minishaker (Supplemental Fig. S1A).

Groups of larvae were subjected to four stimulus intensities (32, 38, 41, and 43 dB re 1 m/s^2^) that were given in increasing order (3 millisecond pulses). The software output frequency was 1000 Hz, though the shaker output of these pulses was more broadband, with variable peaks from 1 to 2000 Hz (Supplemental Fig S1B). Each stimulus intensity was given four to seven times, spaced 20 milliseconds apart to prevent habituation (Table S1) (Wolman et al., 2011). For a given stimulus intensity, startle kinematics were shown to be statistically indistinguishable between the 1^st^ and the 4^th^ stimulus given, indicating that the 20 millisecond wait period was a sufficient interstimulus interval (Supplemental Fig. S2). A high-speed video camera (Edgertronic, CA) was set at a 10% pretrigger rate to capture 13 frames prior to the stimulus being elicited, while recording larval movements at 1000 frames per second (Panlilio et al., 2020).

### Measuring startle vibration

The startle vibration was measured as described earlier (Panlilio et al., 2020). Briefly, vibration was measured using a 3-axis accelerometer (PCB Piezotronics, W356B11). The accelerometer signal was low-pass filtered (10 kHz low-pass cutoff frequency) and amplified with a 30 dB gain before being digitized. Data were converted into acceleration units (m/s^2^) using manufacturer sensitivity values for each axis of the accelerometer and accounting for gain. Data were then band-pass filtered between 1 and 5000 Hz using an 8th order Butterworth filter. The Euclidian norm (vector sum) for the three acceleration signals was calculated to get the total 3D magnitude of acceleration. The maximum value (peak) of each impulse was calculated in decibels (dB) using the following equation:

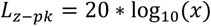

where *L_z-pk_* is the zero-to-peak acceleration level in dB re 1 m/s^2^, and *x* is the maximum acceleration level.

### Direct electric field stimulation

Larvae were head mounted in 35mm glass bottom dishes that were modified by gluing jumper wires to the edges of the dish to create a defined surface area for the electric field. Larvae were positioned rostral-caudally to the electrodes to provide the highest probability of eliciting a startle (Tabor et al., 2014). To head-mount, fish were anesthetized in MS222 (0.16%) and mounted upright (ventral side on the dish) using 1.5% low-melt agarose. Once the agarose solidified, an insect pin (size #000) was used to carefully carve out the agarose so that the region below the head was free to move. The dish was then flooded with embryo media (0.3x Danieau’s) and the fish were allowed to recover in a water bath heated to approximately 26°C. Using an “embryo poker” (a piece of 0.41mm fishing line glued to a glass pipette tip), the fish’s tail was brushed lightly to confirm that it recovered from anesthesia prior to beginning the trial. A 4.4 V/cm, 3 millisecond square wave was generated by the stimulus generator, PulsePal (Sansworks, NY). This was delivered simultaneously as a TTL pulse that triggered the high-speed camera (Edgertronic, CA), which recorded the tail movements at 1000 frames per second. Tail bend angle and latency were then tracked using the Flote software (Burgess and Granato, 2007a).

### Startle behavioral analysis

High speed videos were converted into jpeg files (.mov files with a minimal resolution of 720×720, 1/1008 shutter speed and a frame rate of 1000 frames/second). To reduce the noise and tracking errors, the background was subtracted using a custom script in MATLAB (Panlilio et al., 2020). Flote software (Burgess and Granato, 2007a) was then used to analyze the jpeg files. Quantitative attributes of the startle response measured include startle responsiveness (whether larvae responded or not), latency (time between stimulus and startle), bend angle and maximal angular velocity during startle (Panlilio et al., 2020).

### Using mixture models to identify latency cut-offs for SLC versus LLC responses

In response to auditory/vibrational stimuli, zebrafish perform two types of startle responses: short latency c-bends (SLCs) and long latency c-bends (LLCs) (Supplemental Fig S1C). These two types of startle responses have different latencies, distinct kinematics, and are driven by separate underlying neural circuits (Burgess and Granato, 2007a; Burgess and Granato, 2007b; Higashijima et al., 2003; Marquart et al., 2019). To distinguish between the two types of startle responses, we identified a latency cutoff using a Gaussian mixture model, which fitted two Gaussian distributions, and assigned each latency data point a probability of belonging to either of the two distributions (R package, mixtools) (Benaglia et al., 2009; Panlilio et al., 2020). The cut-off for assigning a response as an SLC was 14 milliseconds - the latency with a greater than 50% probability of belonging to the first fitted Gaussian distribution (Supplemental Fig. S1D). Startle responses that had latencies greater than 14 milliseconds were classified as LLCs.

### Statistical modeling of startle responsiveness

Every fish was given 4-7 replicate auditory/vibrational stimuli per each stimulus intensity tested. For all instances where a fish was successfully tracked, response rates were recorded. Response rates for individual fish were calculated (% responsiveness = number of times the fish responded / number of successfully tracked videos with a maximum of 4 tracks per stimulus intensity per individual fish). A mixed effects logistic regression model was then used to identify treatment differences in percent responsiveness, with the treatment as a fixed effect and the replicate experimental trials as a random effect (glmer(), lme4 R package) (Bates et al., 2015).

### Imaging myelin sheaths in the spinal cord following domoic acid exposures

Our previous results (Panlilio et al., 2020) showed pronounced myelin defects following exposures to DomA at 2 dpf. To determine whether there is a relationship between the behavioral findings and the myelin phenotype, *Tg*(*mbp:EGFP-CAAX*) embryos were exposed to DomA at 2 dpf and screened for myelin defects prior to behavioral analyses at 7 dpf. The severity of the myelin defect was qualitatively scored (Supplemental Fig. S3). To simplify the analyses, four categories were used (Panlilio et al., 2020): (0) Normal phenotype — dorsal and ventral regions had labeled myelin sheaths. The myelin sheath surrounding the Mauthner axon was visible. (1) Myelin sheaths were present but disorganized. In some cases, myelinated axons that were normally found ventrally were located more dorsally. In others, the myelinated axons terminated prematurely with distal ends located more dorsally. (2) Myelin was labeled in both the dorsal and ventral regions of the spinal cord, but there are some noticeable deficits. While the ventral spinal cord was labeled, it had noticeably less myelin labeled compared to controls. (3) The loss of labeled myelin in the ventral spinal cord resulted in large, observable gaps between myelinated axons. Circular myelin membranes were present both in the ventral and dorsal spinal cord.

### Statistical modelling of the distribution of startle response type due to treatment

When fish responded to a stimulus, they either performed an SLC or an LLC startle which was defined by the latency cut-off of 14 milliseconds. To determine whether domoic acid exposure alters the proportion of SLC to LLC responses, a mixed effects logistic regression model was used with treatment as a fixed effect and the replicate experimental trials as a random effect.

DomA-exposed fish were also classified by the severity of myelin defects. To determine whether there is an association between the myelin severity and the likelihood that a fish performs an SLC rather than an LLC, a separate mixed effects logistic regression model was used, with myelin severity as the fixed effect and the replicate experimental trials as a random effect.

### Calculating the Startle Bias Index

The Startle Bias Index was calculated as described previously (Jain et al., 2018). Bias per individual fish was calculated as the (frequency of SLC – frequency of LLC)/ total responses where +1 represents the value in which all responses were SLC startles, and −1 represents the value in which all were LLC startles. Following this, mean behavioral biases for a treatment group were calculated.

### Analysis of treatment differences in startle response kinematics (Panlilio et al., 2020)

Kinematic responses from the two types of startle responses (SLC v. LLC) were analyzed separately based on previous research showing that they are driven by distinct neural circuits and have distinct kinematic characteristics (Burgess and Granato, 2007b; Marsden and Granato, 2015; O’Malley et al., 1996). The median response of individual fish for each startle type was then calculated for each intensity level.

To determine whether DomA alters kinematic responses to startle, we performed Aligned Ranked Transformed ANOVA test, with treatment group as a fixed factor (DomA vs. control). Nine separate repeated experiments (trials) were combined for the analysis (art(), ARTool R package) (Wobbrock et al., 2011) (Table S1). To account for potential differences in response due to the variations between trials, the experimental trial was incorporated into the model as a random factor.

To determine whether there was an association between the myelin phenotypes and the startle kinematics, we used a nonparametric multiple comparisons test with Dunnett-type intervals (nparcomp(), nparcomp R package) (Konietschke et al., 2015).

### Immunohistochemistry

Fish were fixed in 4% paraformaldehyde overnight at 4°C. In some cases, whole brains were dissected, and in other cases whole embryos were used. Antigen retrieval was done by placing tissue in 150 mM Tris HCl (pH 9.0) in a 70°C water bath for 15 minutes (Inoue and Wittbrodt, 2011). Brain tissue was permeabilized using proteinase K (10 μg/ml for 3 minutes) then post-fixed for 20 minutes using 4% paraformaldehyde. Whole mount embryos were permeabilized using ice-cold acetone (7 minutes). Samples were then blocked in 10% normal goat serum and 1% DMSO, followed by 1-3 day incubations in primary antibodies (α-3A10 −1:100 dilution, Developmental Studies Hybridoma Bank, antibody registry ID: AB 531874, prepared 7/28/16, 51 μg/ml; α-acetylated tubulin - 1:500 dilution, Santa Cruz Biotechnology, SC 23950 sample HS417, 10 μg/ 50 ml). After several washes in phosphate buffered saline with 0.01% triton-X, samples were then incubated in secondary antibodies (1:400 Alexa Fluor 488 Goat α-mouse, Catalog A11001, 1752514 or Alexa Fluor 594 Goat α-mouse; Abcam, GR196875). Samples were then placed in antifade mountant (Prolong or SlowFade Diamond mountant, Invitrogen), and placed between bridged #1.5 coverslips for imaging.

Acetylated tubulin was used to assess primary neuron axons, early sensory neurons (Rohon beard cells), and the peripheral lateral line, while 3A10 was used to characterize Mauthner cell bodies and hindbrain and midbrain axonal tracks. Brain dissections required for 3A10 staining sometimes led to reticulospinal axons (those that extend down to the spinal cord) being removed from the preparation. (Supplemental Fig. S4 shows the range of phenotypes found in controls). Thus, for the antibody labeling experiment, the presence or absence of the Mauthner cell was determined solely based on the presence of the Mauthner cell body and lateral dendrite.

### DASPEI labeling

Sensory neuromasts were labeled using the vital dye, DASPEI ((2-(4-(dimethylamino)styryl)-N-ethylpyridinium iodide, Biotium Inc., Freemont, CA). Zebrafish larvae (5 dpf) were incubated in 0.005% DASPEI for 15-20 minutes, then washed and placed in anesthetic (0.16% MS222). Larvae were then mounted in low-melt agarose (1-1.5%) either laterally or ventrally and imaged with a widefield microscope. Neuromasts were manually counted on blinded files. To identify treatment differences in counts, a random coefficient Poisson regression was done, with repeated experiments (trials) modeled as a random factor (glmer(), lme4 R package) (Bates et al., 2015).

### Reticulospinal backfills

Larvae (7 dpf) were anesthetized using MS222 (0.16%), then mounted ventrally in 1.5% low melt agarose. Dissection spring scissors were dipped into Texas red dextran (3000 MW) and the spinal cord was transected at the level of the anus (Moens et al., 1996). Larvae were placed back into anesthetic and screened for dye uptake using epifluorescence microscopy (Zeiss Axiovert inverted microscope). Roughly an hour after spinal cord transections, larvae were placed in 4% paraformaldehyde and fixed overnight in 4°C. Following several washes in PBS, the whole brain was dissected (Turner et al., 2014). Whole brains were mounted using the same technique used with brains that were labeled with antibodies. The presence of the Mauthner cell, along with any labeled neurons in rhombomere 5 (r5) and rhombomere 6 (r6) - the location of MiDcm2, MiDcm3 – was identified. Following this classification, ordered logistic regression was done to determine whether treatment alters the number of neurons found in each location (polr(), Mass R package, R) (Ripley et al., 2018).

### Primary motor neuron live imaging and tracing

*Tg*(*cntn1b:EGFP-CAAX*) embryos were exposed to DomA at 48-53 hpf. Larvae (2.5 dpf) were then anesthetized (0.16% MS222), and embedded laterally in 1-1.5% low-melt agarose in glass bottom microscopy dishes. Larvae were imaged using a confocal microscope (LSM 710 or LSM 780) with the 40x water immersion objective (C-Apochromat, 40x, NA 1.1). Primary motor neurons located approximately at myotome 20-23 were imaged. Images were blinded prior to image analysis. A single primary motor neuron from the image stack was chosen. Its main axon along with the axon collaterals were traced from the image stacks using the ImageJ plugin, Simple Neurite Tracer (Longair et al., 2011). The total length of the traced axon collaterals was calculated from the tracing files. One-way analysis of variance was used to determine whether there was an effect of treatment on the total length of the axon collaterals (oneway.test(), R).

## RESULTS

### DomA-exposed fish were less responsive, and less likely to perform SLC startles

We first determined whether DomA alters responsiveness over the range of stimulus intensities tested. At the lowest stimulus intensity (32 dB), most control and DomA-treated fish did not respond (Fig. 1A, Table S2). Furthermore, both DomA and control fish that did respond were more likely to perform LLCs rather than SLC startles. While this was true for both control and DomA-exposed fish, DomA-exposed fish were even less likely to respond (Estimate = −0.474, p=0.0132, Table S3), and even less likely to perform SLC startles rather than LLC startles compared to controls (Estimate = −1.076, p=5.84E-15, Table S4).

**Figure 1:**
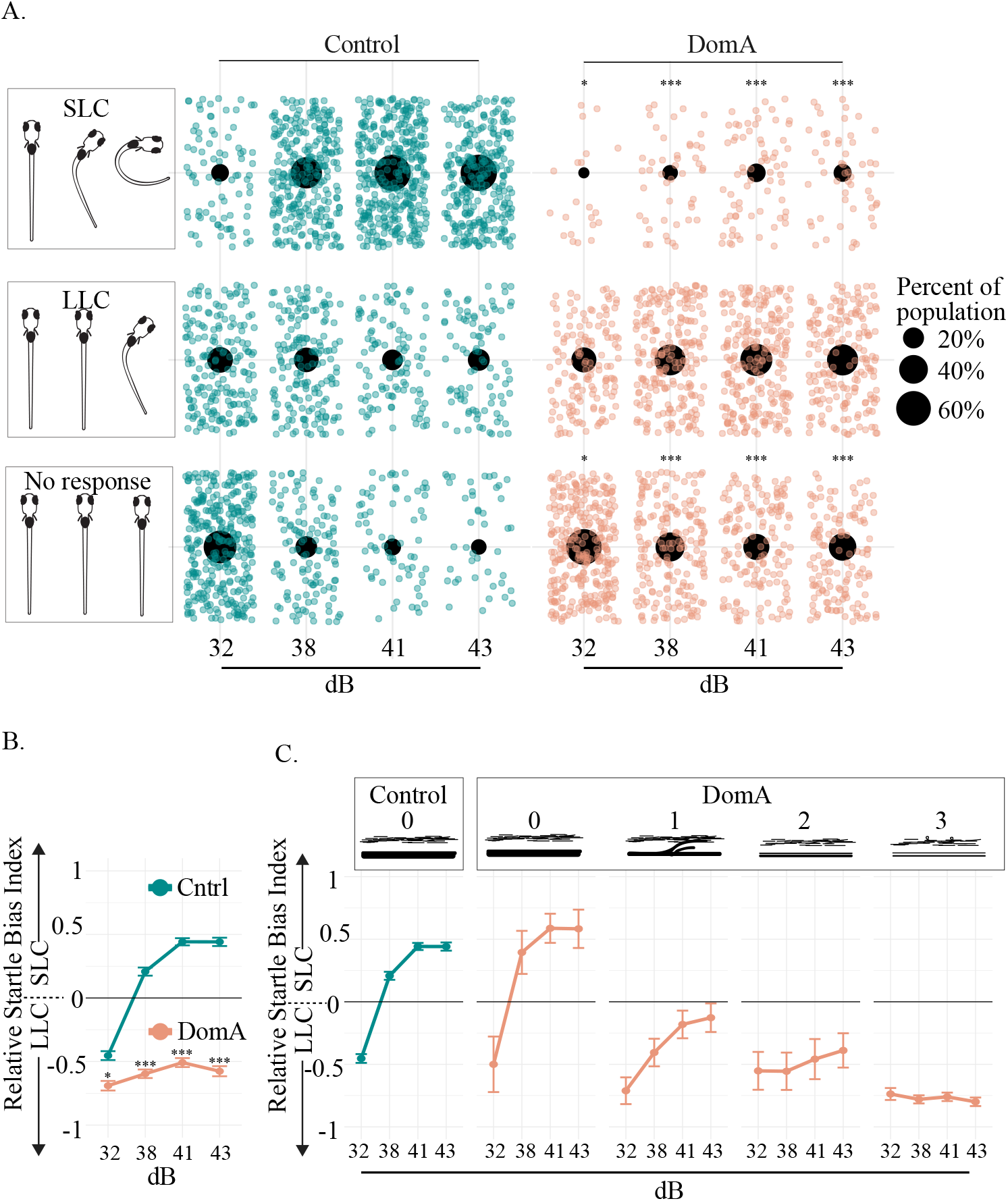
Domoic acid-exposed larvae are less responsive and preferentially perform LLC startles compared to controls when given auditory/vibrational stimuli. **(A)** Distribution of fish that did one of three behaviors: 1) no response, 2) LLC startle, or 3) SLC startle for each stimulus intensity. Only the first of the 4 replicate stimuli were graphed so that each fish is represented once. Individual points represent single fish that performed each behavior. Large black dots represent the proportion of the population that performed one of the three behaviors within a given stimulus intensity. Asterisks in the no response column represent statistically significant differences in responsiveness in DomA-exposed fish relative to controls within each stimulus intensity. Asterisks in the SLC column represent statistically significant differences in the performance of SLC rather than LLCs in DomA-exposed fish relative to controls within each stimulus intensity. Significance was determined using mixed effects logistic regression. *p < 0.05, *** p< 0.001 **(B)** Relative Startle Bias Index was calculated for all fish that were responsive. Individual fish were provided with 4 replicate stimuli within a given stimulus intensity. Bias per individual was calculated as the (frequency of SLC - frequency of LLC)/ total responses. +1 represents the value in which all responses were SLC-type startles, and −1 represents the value in which all were LLC-type startles. Mean behavioral biases for a treatment group per stimulus intensity were graphed. Asterisks indicate statistical significance in performing SLC startles (versus LLC startles). **(C)** Domoic acid-treated larvae were also classified by myelin category (0-3) in a subset of the experiments (subset of fish graphed in Figure 1B). Startle bias per myelin phenotype was plotted.

With higher stimulus intensities (≥38 dB), a greater proportion of both the control and DomA-treated fish performed startles rather than not responding (Fig. 1A, Table S2). However, as seen at the lowest stimulus intensity, DomA-exposed fish were significantly less likely to respond and significantly less likely to perform an SLC startle vs. a LLC startle, compared to control fish (Tables S3 and S4).

SLC loss is further illustrated by calculating the *startle bias index* (Jain et al., 2018), a measure of the frequency with which individual fish performed an LLC versus an SLC response at a given stimulus intensity. At intensities of 38 dB and higher, control fish preferentially performed SLC rather than LLC responses (Fig. 1B). In contrast, DomA-exposed fish preferentially performed LLC rather than SLC responses at all stimulus intensities tested.

### Fish with myelin defects were less likely to perform SLC startles than those without defects

Previously, we showed the DomA-exposed larvae had myelin deficits (Panlilio et al., 2020). To assess the relationship between startle bias and the observed myelin sheath defects, we classified a subset of the larvae based on myelin defects prior to the behavioral assessment. Similar to the controls, DomA-exposed fish with normal myelin sheaths (category 0) performed more SLC startles at higher stimulus intensities (38 dB and higher) and did not differ from controls in the likelihood of performing LLCs over SLCs at the highest intensity tested (43 dB) (Fig. 1C, Table S5). In contrast, fish with any noticeable myelin defects (Category 1-3) had a reduced likelihood of switching from LLC to SLC startles with increasing stimulus intensity, as compared to controls (p <0.0004, Table S5). Furthermore, DomA-exposed fish with the most severe defects (Category 3) were also less likely to switch from LLC to SLC startles at 43 dB compared with DomA-exposed fish with any of the less severe myelin defects (Category 0-2) (Estimate= −3.196 for DomA-exposed fish (Category 3) = −3.1957, p= 4.81E-14; Table S5).

### DomA-exposed fish that had myelin defects also had startle response kinematic deficits

For the fish that responded to a given stimulus, the startle kinematics (bend angle, maximal angular velocity) were assessed. In agreement with our previous findings (Panlilio et al., 2020), DomA-exposed animals had smaller bend angles and lower maximal angular velocities (Mav) compared to control animals at all the intensities tested (Supplemental Fig. 5; p < 2.22 e-16). This was true for both LLC (Supplemental Fig. 5A, B; Table S6) and SLC (Supplemental Fig. 5C, D) startles.

We then determined whether myelin defect severity was correlated with startle kinematic performance. A majority of DomA-exposed larvae were classified as category 3 (the most severe myelin defects). When performing either SLC or LLC startle responses, these larvae had significantly reduced bend angles and Mavs for all stimulus intensities tested (Fig. 2).

Fish with intermediate myelin defects (category 1-2) also consistently had kinematic deficits when performing SLC startles across all stimulus intensities tested (Fig. 2A, 2B, Table S6). However, it was only fish with the most severe myelin defects (category 3) that had reduced bend angles and slower maximal angular velocities during LLC startles (Fig. 2C, 2D, Table S7).

**Figure 2:**
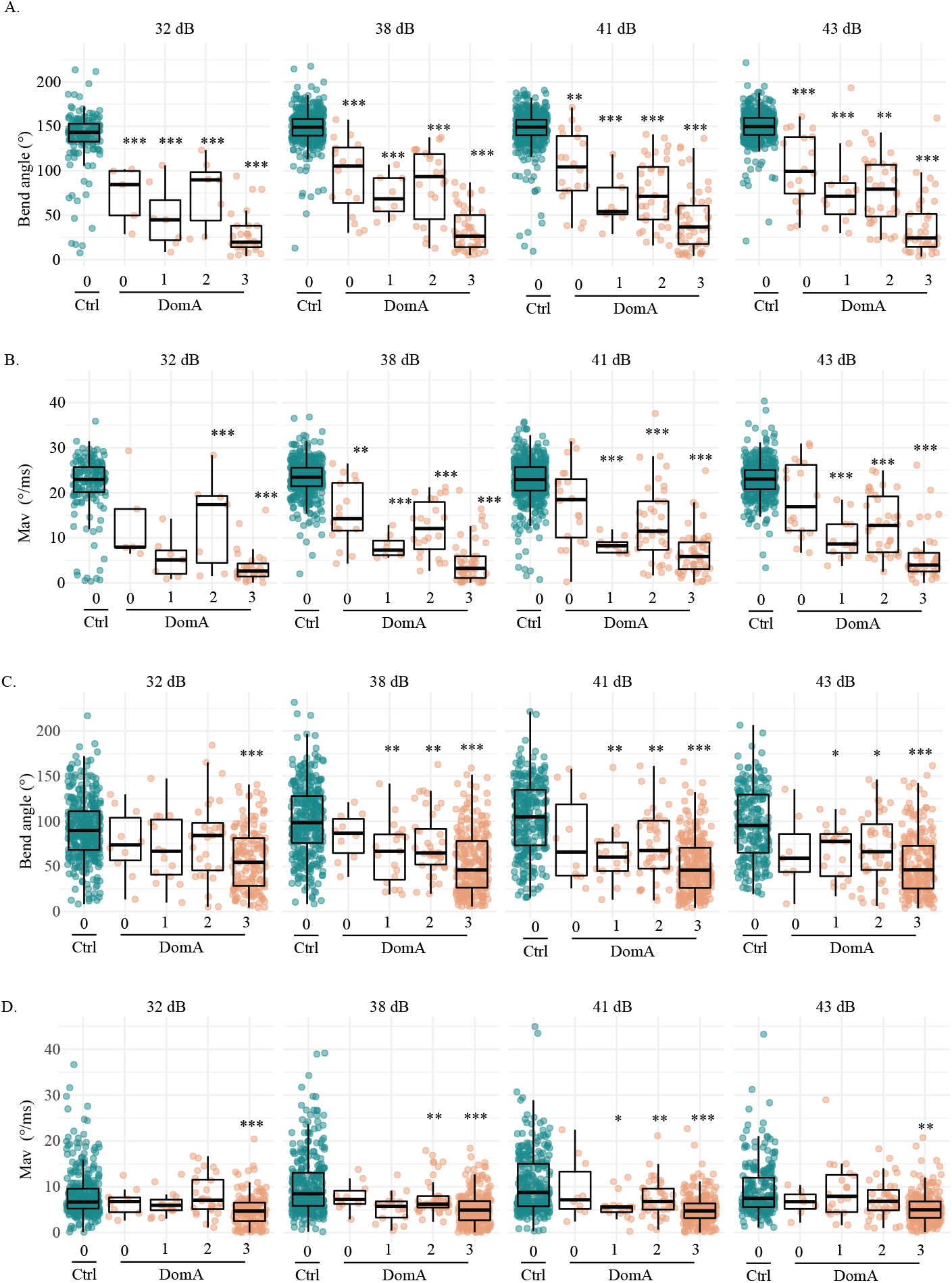
Kinematics deficits in auditory/vibrational startle are correlated to myelin defects. **(A)** DomA-treated fish were further subcategorized by myelin sheath defects. 0 = control-like myelin sheaths to 3 = the most severe myelin defect observed. Bend angles during SLC startles with increasing stimulus intensities. **(B)** Maximal angular velocities (Mav) during SLC startles with increasing stimulus intensities. **(C)** Bend angles during LLC startles with increasing stimulus intensities. **(D)** Maximal angular velocities (Mav) during LLC startles with increasing stimulus intensities. Asterisks represent statistical significance between DomA and controls determined using an Aligned Ranked Transformed ANOVA test (* = p <0.05, ** = p< 0.001, *** = p < 0.0001).

Like DomA-treated fish in other categories, DomA-treated fish with control-like myelin sheaths (category 0) also had kinematic deficits when performing SLC startle responses (Fig. 2A, 2B, Table S6). In contrast, DomA-treated fish with category 0 phenotypes did not have any measurable kinematic deficits when performing LLC startle responses at all stimulus intensities tested (Fig. 2C, 2D, Table S7).

These results suggest that exposures to DomA that led to distinguishable myelin defects also led to startle deficits. The severity of the myelin defect was correlated with the severity of the behavioral deficit; fish with the most severe myelin defects were also less likely to perform SLCs and had shallower bend angles and slower maximal angular velocities relative to controls.

### DomA-exposed fish were less responsive, and had aberrant kinematics following direct-electric field stimulation

To assess the contribution of the sensory system to the observed startle deficits, we used direct electric field stimulation, which bypasses the sensory system, directly activating the Mauthner cell, leading to ultra-rapid startle responses (Tabor et al., 2014) (Fig. 3A). DomA-treated larvae were significantly less responsive to electrical stimulation than control fish (p < 1 e-16). A majority of control fish (70/74) responded to all of the 7 replicate stimuli (Fig. 3B), while a majority of the DomA-treated fish (47/74) did not respond to any of the replicate stimuli. Furthermore, control fish responded rapidly to electrical stimulation, with a median latency of 2 milliseconds (Fig. 3C). Strikingly, when DomA-treated fish did respond, they had significantly longer latencies (control median = 2 ms, DomA median= 49 ms, IQR= 2 s, estimated relative effect (est) = 0.78, 95% CI [0.633, 0.927], p=0.001). When DomA-treated fish did respond, they also had shallower bend angles and slower maximal angular velocities compared to controls (bend angle: median =83.5° in controls, median = 41.9° in DomA-treated, Coefficient= 0.173, 95% CI [0.059, 0.286], p < e −16; mav: median = 3.78 °/ms in controls, median = 2.28 °/ms in DomA treated, Coefficient = 0.21, 95% CI [0.083, 0.338], p < e −16) (Fig. 3D and 3E).

**Figure 3:**
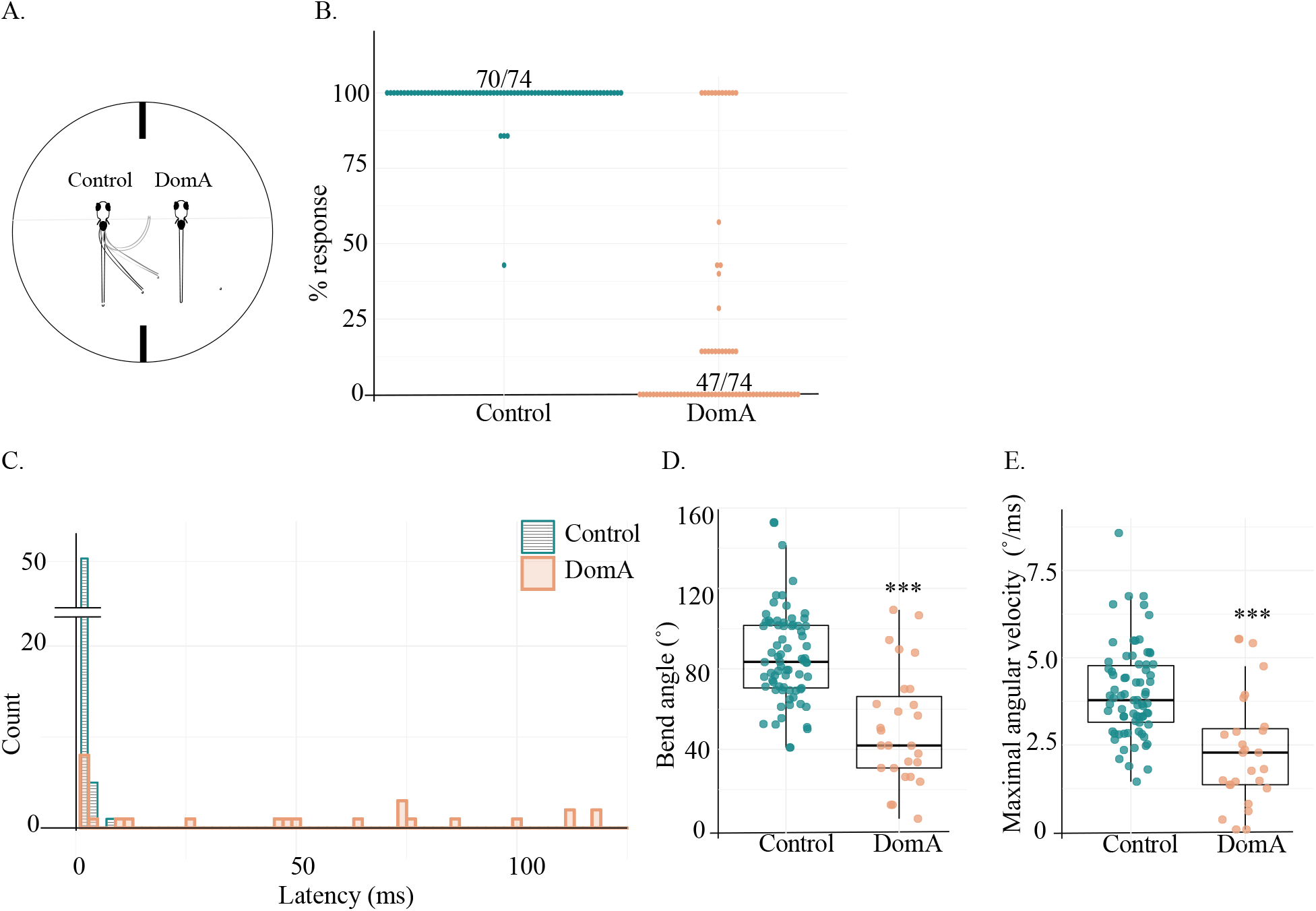
Domoic acid-exposed larvae have aberrant startle responses to direct electrical stimulation. **(A)** Larvae were head-mounted in agar and positioned rostral-caudally to the electrodes. Control larvae were mounted on the left, DomA-exposed larvae on the right. Larvae were then provided with a 6 Volt, 2 millisecond (ms) square pulse. Their tail movements were captured using a high-speed video camera. **(B)** Percent responsiveness of individual larvae to 7 identical electric field pulses. Points represent the percent of times an individual fish responded to replicate stimuli. 70/74 of the control population responded 100% of the time while 47/74 of DomA-treated population responded 0% of the time. **(C)** Latency distributions for control and DomA-treated fish. Count represents the number of fish that have latencies within each of the 2 ms time bins. The majority of the control fish responded within 2 ms after stimulus was produced (51/74 Control fish had a median latency of 1-2 ms). Not shown= 1 DomA-treated larva that responded at 179 ms. **(D)** Maximal bend angle for control (n= 74) versus DomA-treated fish (n= 27). Each point represents the median response of an individual fish. **(E)** Maximal angular velocity for control versus DomA-treated fish. Each point represents the median response of an individual fish. ***p ≤ 0.0001 indicates significant difference between DomA-exposed fish relative to control fish. Statistical significance was determined using a nonparametric Behrens-Fisher t-test.

Like auditory/vibrational startle deficits, deficits in electric field-induced startle were also correlated with myelin defects; DomA treated fish with more severe myelin defects also had more deficits in startle (Fig. 4). DomA-treated fish that had no visible myelin defects had no differences in responsiveness, bend angles, Mavs, or latencies compared to controls (Fig. 4A-E). In contrast, DomA-treated fish with any visible myelin defect, even in its least severe form (category 1), had significantly reduced responsiveness compared to controls (Dunnet post-hoc test, p < e −9) (Fig. 4E). Furthermore, DomA-exposed fish that had defects of intermediate severity (category 2) also had significantly longer latencies (est= 0.989, 95% CI= [0.946, 1.032], p<3.28 e −13), but no statistically significant differences in bend angle or Mav (Fig 4C, 4D; Table S8). Finally, DomA-exposed fish that had the most severe myelin defects (category 3) had deficits for all startle attributes measured (Fig. 4A-E; Table S8).

**Figure 4:**
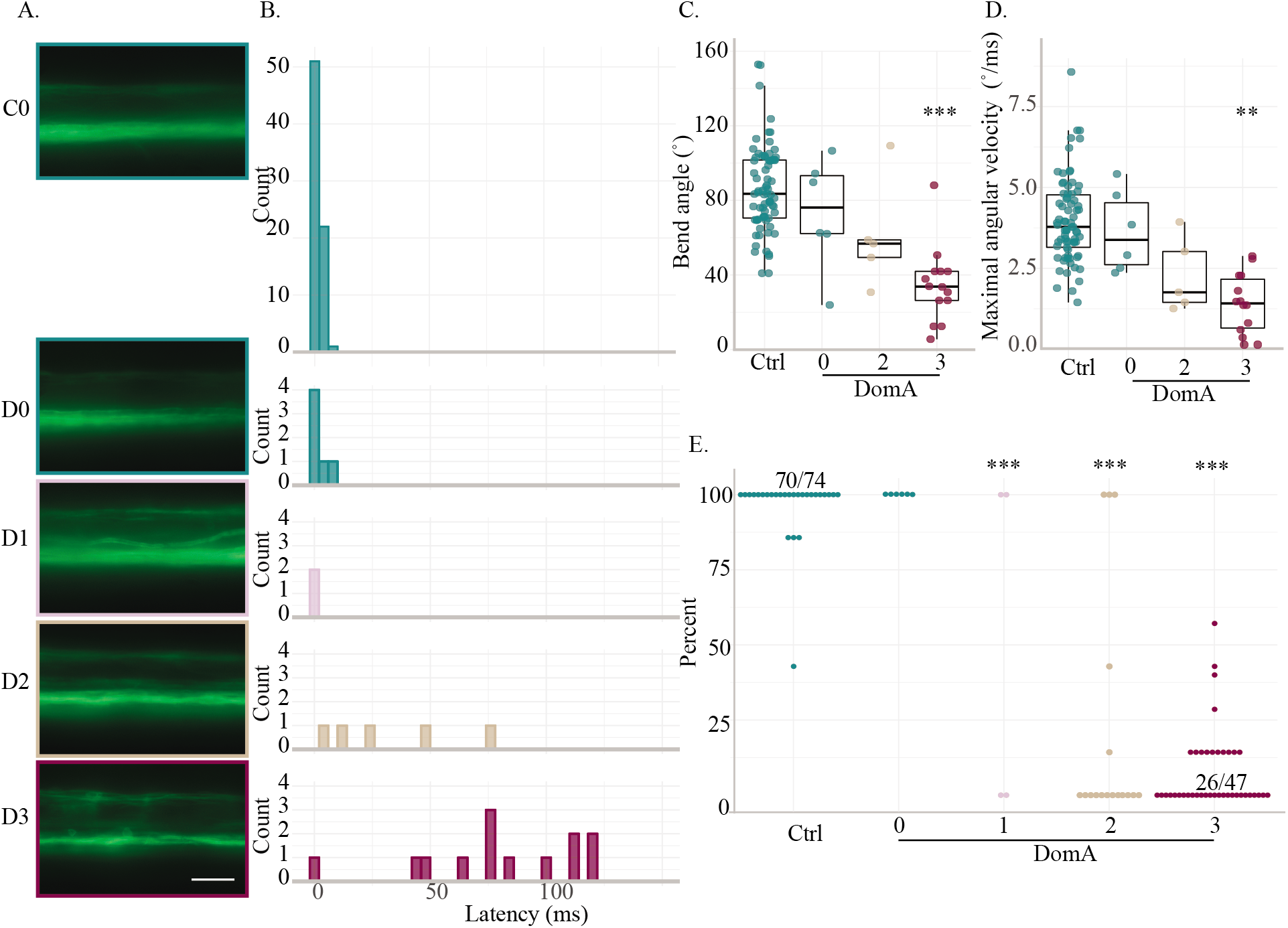
Severity in myelin defects is correlated with startle kinematic deficits from direct electric field stimulation. **(A)** Representative widefield epifluoresence images of the myelin sheath phenotypes categorized, with (0) having no myelin defect to (3) having the most myelin severe defect observed. C0 = Control, D0-D3= DomA treated. **(B)** Latency distributions for control and DomA-treated fish. The same data used in Figure 5C but further categorized by myelin sheath imaging categories. Not shown= 1 DomA-treated larvae which responded at 179 ms. **(C)** Maximum bend angle for control (n=74) versus DomA-treated fish (n=25). Fish with myelin defects in category 1 were excluded from this analysis due to low sample sizes (n=2). **(D)** Maximal angular velocity for control versus DomA-treated fish. **(E)** Percent responsiveness of individual larvae to 7 identical electric field pulses. Points represent the percent of times an individual fish responded to replicate stimuli. Ratios listed above a group of points represent the ratio of fish that responded to that stimulus over the total within that percentage bracket. Ratios were listed when all the individual points could not fit on the graph. Asterisks represent statistical significance between DomA and controls determined using the a nonparametric multiple comparisons test with Dunnett-type intervals *** = p < 0.0001).

To further identify the potential cellular and structural defects underlying these observed behavioral deficits, we imaged different components of the startle circuit, starting with the sensory system.

### Sensory inputs for acoustic/vibrational startle

We assessed the key sensory inputs that are required for performing auditory/vibrational startle responses. When an auditory/vibrational stimulus is provided, hair cells in the inner ear, and in some circumstances within neuromasts in the lateral line, are activated. When activated, these hair cells lead to the activation of the statoacoustic ganglia (in the inner ear) or the lateral line ganglia, which in turn send the sensory information to the hindbrain where the information is integrated (Nicolson, 2017; Nicolson et al., 1998). To assess whether DomA disrupts the sensory system, we sought to determine whether DomA reduced the number of neuromasts or disrupted statoacoustic ganglia or lateral line structures.

Using the vital dye DASPEI, we labelled the neuromasts and found no differences in the number of neuromasts in both the cranial and the trunk region after exposure to DomA (Fig. 5A,B). By exposing transgenic *Tg*(*cntn1b:EGFP-CAAX*) fish, in which the peripheral lateral line and the statoacoustic ganglia are labeled, we also found no discernible differences in the presence of these structures compared to controls (Fig. 5D,E).

**Figure 5:**
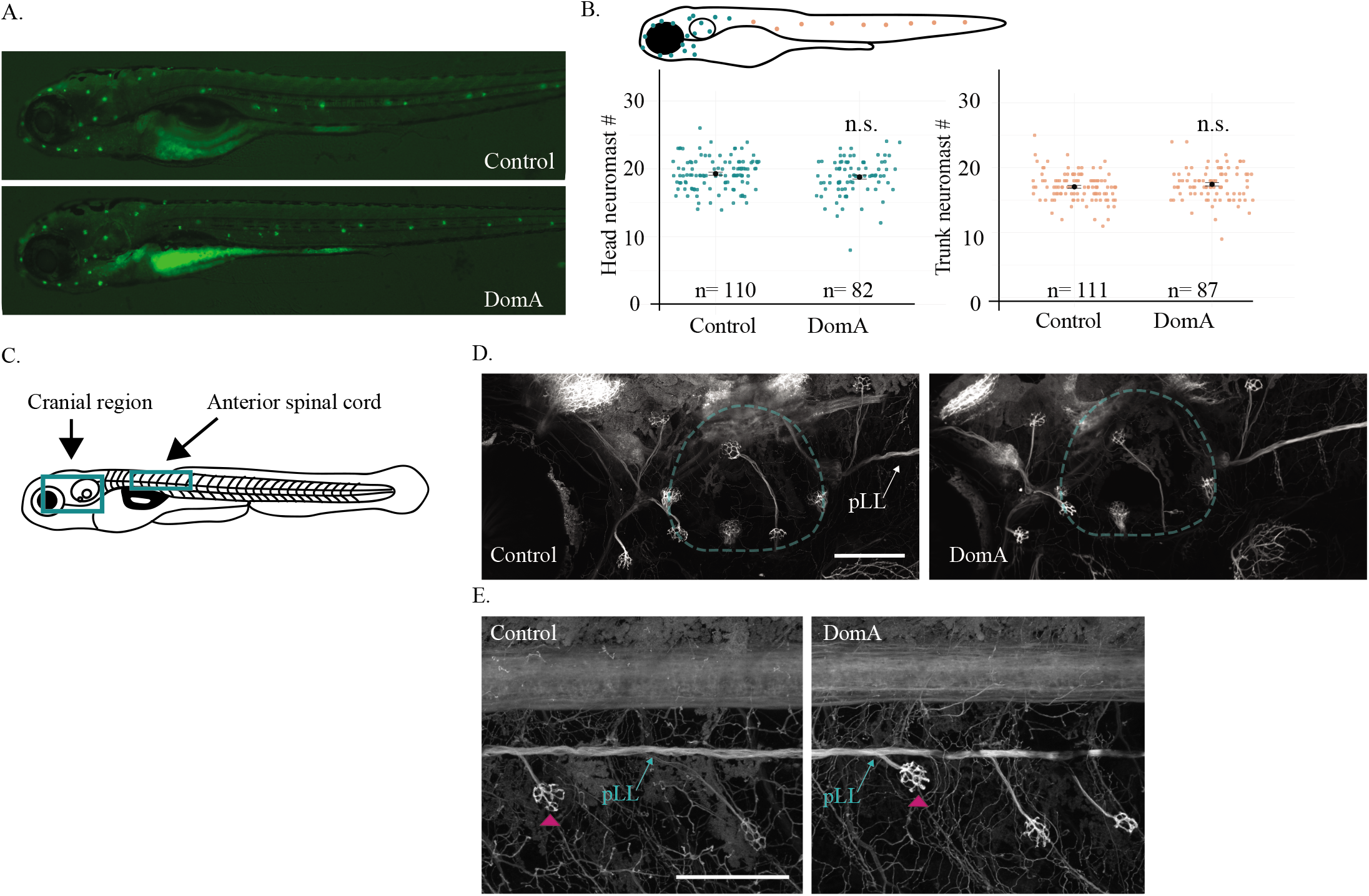
The sensory inputs required for the startle response appeared intact in domoic acid-exposed fish. **(A-B)** DASPEI labeling of sensory neuromasts in 5 dpf larvae. **(A)** Representative widefield fluorescence images of DASPEI-stained control and DomA-exposed larvae. **(B)** Diagram of a 5-dpf larva with head neuromasts colored in teal and trunk neuromasts colored in peach. Head and trunk neuromast counts for control and DomA-exposed. Single points represent individual larvae. Black bar represents standard error of the mean (SE). For cranial region, control - mean = 19 ± 2 (SD), DomA - mean = 19 ± 3 (SD) For trunk region, control - mean =17, ± 2 (SD), DomA - mean = 17 ± 3 (SD) **(C-E)** Imaging of sensory ganglia in 5 dpf *Tg*(*cntn1b:EGFP-CAAX*) larvae. **(C)** Diagram of a laterally mounted *Tg(cntn1b:EGFP-CAAX)* larva, with green boxes that indicate approximate areas imaged. **(D)** Representative confocal images from the cranial region of laterally mounted *Tg*(*cntn1b:EGFP-CAAX*) control (n=33) and DomA-exposed (n=45) larvae. Inner ear is outlined in teal. **(E)** Representative confocal images from anterior spinal cord in laterally mounted *Tg*(*cntn1b:EGFP-CAAX*) control (left; n=36) and DomA-exposed (right; n=46) larvae. Pink arrowhead points to a neuromast. pLL = peripheral lateral line. Scale bars: 100 μm.

### Reticulospinal neurons integrate sensory information

To determine whether DomA disrupts hindbrain neurons responsible for integrating sensory information, we performed spinal backfills. This allowed us to label the Mauthner cells – paired neurons required for eliciting SLC responses - and their homologs MiDcm2 and MiDcm3, which are active during LLC responses (Marsden and Granato, 2015; O’Malley et al., 1996). Most of the control larvae had both Mauthner cells and all four neurons within rhombomere 5 (r5) and rhombomere 6 (r6), the position where MiDcm2 and MiDcm3 reside (Fig. 6). In contrast, a majority of DomA-exposed larvae had neither Mauthner cell, and 0 or 1 (out of the 4) neurons in positions r5 and r6 (Fig. 6A-C). In fact, fish exposed to DomA were 99.3% less likely than controls to have both Mauthner cells rather than 1 or 2 Mauthner cells (odds ratio (OR) = 0.007, p = 1.7 e −14), and had an even lower probability of having 4 neurons in r5 and r6 (OR = 0.001, p= 1.6 e −09).

**Figure 6:**
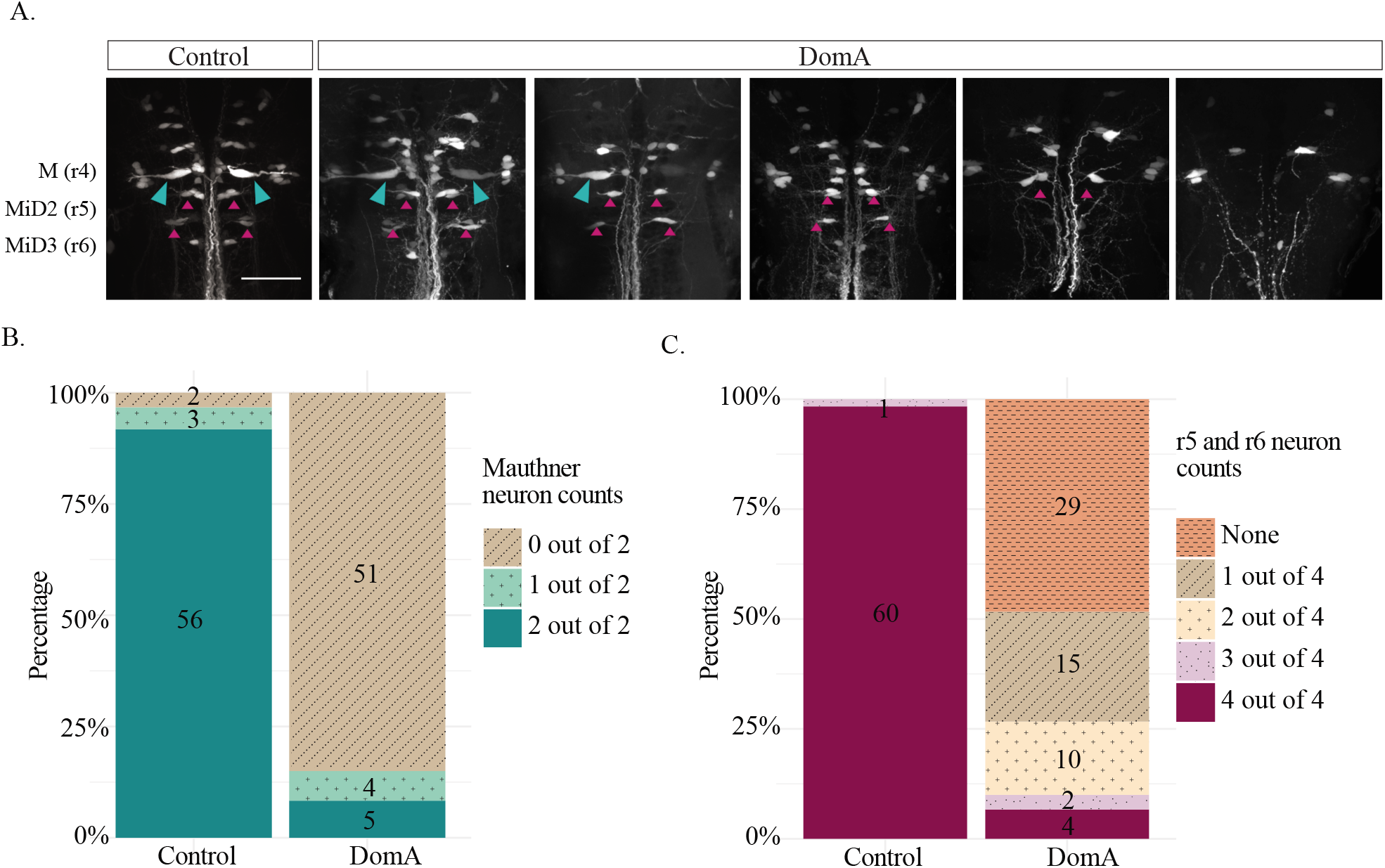
The majority of reticulospinal neurons required for startle responses are absent in domoic acid-exposed larvae. **(A)** 7 dpf larvae were backfilled with Texas Red dextran through spinal cord transections. The figures represent the range of phenotypes observed in control and DomA-injected fish. Teal arrows mark Mauthner cells and magenta arrows mark backfilled neurons in rhombomere 5 (r5) and rhombomere (r6). **(B)** Mauthner cells on the two lateral sides were scored per fish. A majority of DomA-exposed fish (identified 51 out of 60) did not have any Mauthner cells. **(C)** Other reticulospinal neurons involved in startle responses (MiD2cm, MiD3) are located in r5 and r6. The presence of any neuron backfilled in r5 and r6 on the two lateral sides was scored. A majority of DomA-exposed fish had one or no neurons that were backfilled in this r5 and r6 (44 out of 60). Statistical significance was determined using ordered logistic regression. Scale bar = 50 μm.

Spinal backfills have some disadvantages. While the absence of labeled neurons may be an indication that the neurons are not present, it is possible that the neurons are present but may not have axons that extend to the cut site where the axons can take up dye. Furthermore, if neurons have defective axonal transport mechanisms, their axons may be present but unable to take up the dye efficiently enough to stain their cell bodies. To address these potential pitfalls, we also performed whole brain 3A10 antibody staining, which labels the Mauthner cell bodies along with other midbrain and hindbrain axonal tracks (1-7) (Fig. 7A). In agreement with the spinal backfill data, a majority of DomA-exposed larvae did not have Mauthner cell bodies, while a majority of controls did (Fig. 7B and 7C). Importantly, other axonal tracts in the midbrain and hindbrain appeared to be intact in both control and exposed animals. Both control and DomA-exposed animals had medial longitudinal fasciculi, and there were no significant differences in the number of hindbrain axonal tracts (Fig. 7B and 7D).

**Figure 7:**
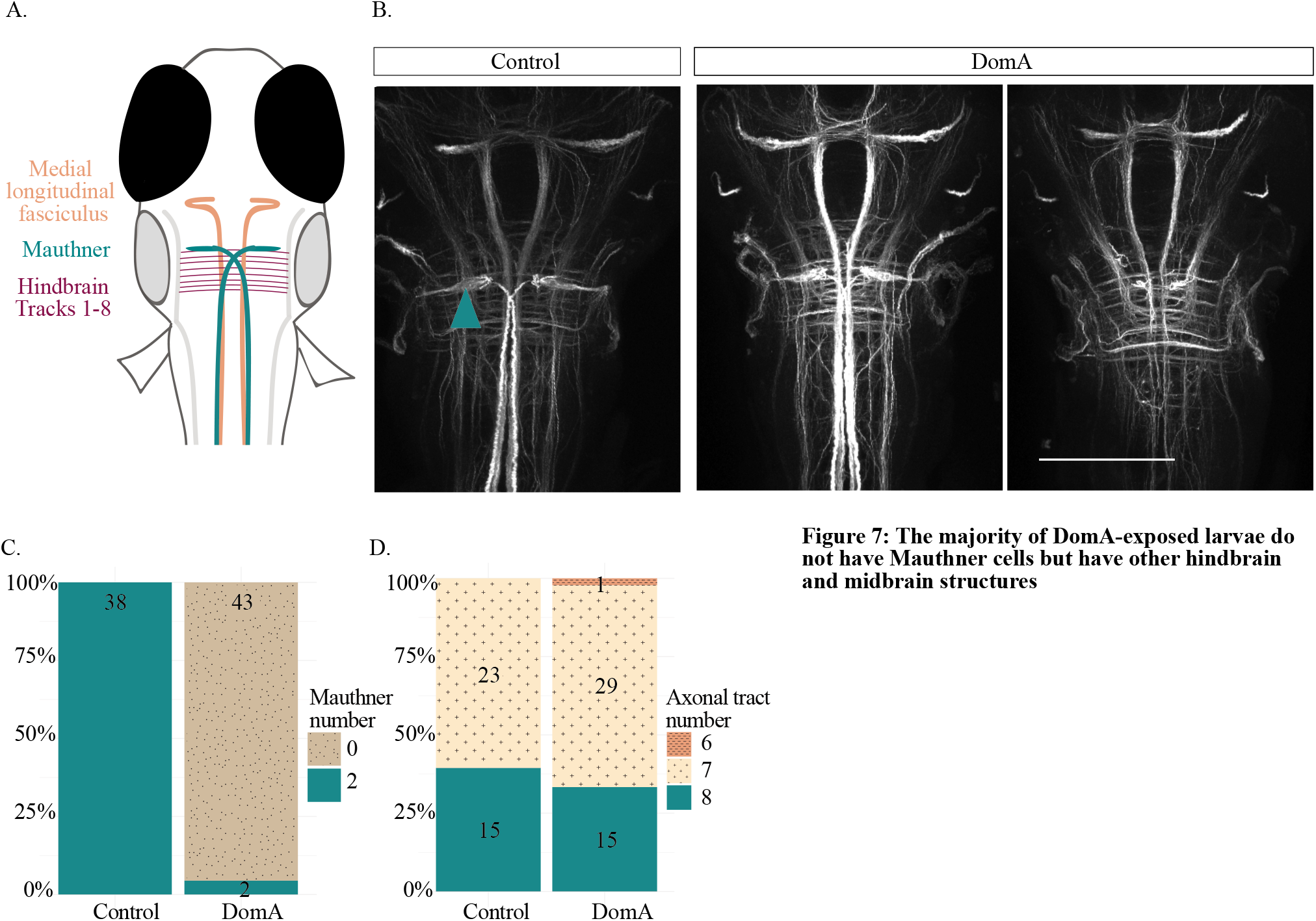
The majority of DomA-exposed larvae do not have Mauthner cells but have other hindbrain and midbrain structures. **(A)** Schematic of brains stained with 3A10, with labels for the Mauthner cells, medial longitudinal fasciculus, and hindbrain axonal tracts. **(B)** Representative images of control and DomA-exposed brain tissue stained with 3A10. Teal arrow points to Mauthner cell. Scale bar = 100 microns **(C)** Score of the number of Mauthner cells (0-2) present in control and treated larvae. Numbers within each section represent the number of larvae with the given phenotype. **(D)** Score of the number of hindbrain axonal tracts detectable in control and treated larvae. Numbers within each section represent the number of larvae with the given phenotype.

### Primary motor neurons that innervate muscles that generate the response

We then determined whether DomA disrupts primary motor neurons that synapse with the hindbrain reticulospinal neurons and innervate the muscles required to elicit startle. To determine whether DomA alters the main primary motor neuron axons, we stained the trunk region using α-acetylated tubulin (Fig. 8A). There were no differences in the presence of the primary motor neurons in DomA-exposed larvae versus controls (Fig. 8B,C). In the same tissue, there were also no differences in the appearance of sensory neuron cell bodies or the peripheral lateral line (Fig. 8B,C).

**Figure 8:**
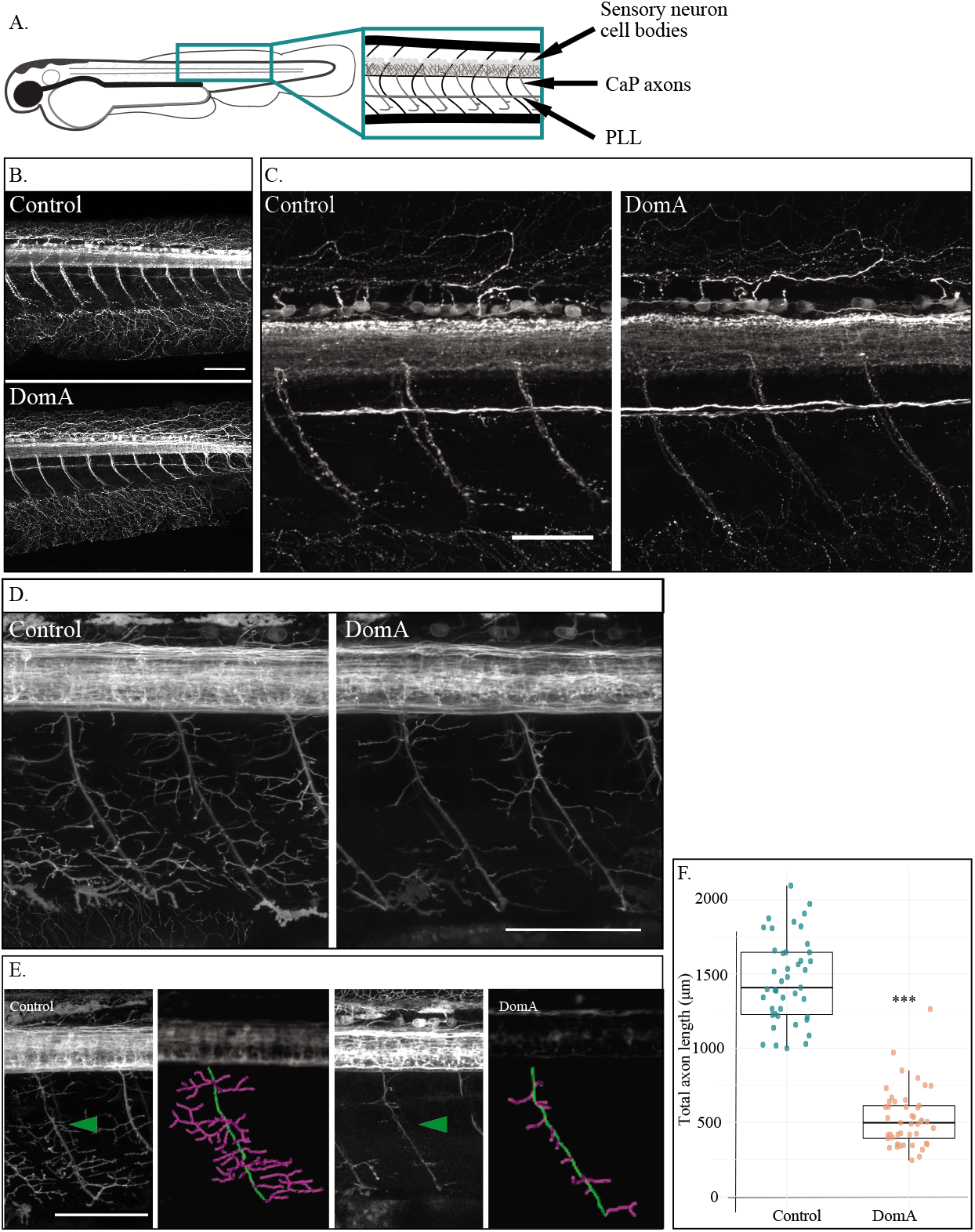
Domoic acid exposure reduces axon collateral branching in primary motor neurons. **(A-B)** Immunostaining for acetylated tubulin in 2.5 dpf embryos **(A)** Diagram of a 2.5 dpf larvae laterally mounted and immunostained with α-acetylated tubulin. Anatomical features of interest are highlighted (PLL= peripheral lateral line, CaP= caudal primary motor neurons). **(B)** Immunostaining with α-acetylated tubulin at 2.5 dpf. DomA-exposed embryos (n=34) had visible CaP axons, sensory neuron cell bodies, and PLLs that were indistinguishable from controls (n=31). **(C)** Higher resolution image of control and DomA-exposed fish immunostained with α-acetylated tubulin **(C-D)** Imaging of primary motor neuron branching in 2.5 dpf *Tg(cntn1b:EGFP-CAAX)* embryos **(D)** Representative images from 5 dpf *Tg*(*cntn1b:EGFP-CAAX*) controls and DomA-exposed larvae taken at high resolution. **(E)** Representative images from 5 dpf *Tg*(*cntn1b:EGFP-CAAX*) controls and DomA-exposed larvae used for tracing studies. Green arrow points to motor neuron axons that were traced to estimate primary motor neuron axonal lengths in each treatment. Tracings are false colored. The main primary motor neuron axon was traced in green, and the axon collaterals were traced with magenta. **(F)** Quantification of total lengths from axonal tracings of a subset of the imaged fish (inclusive of the main motor neuron in green and axon collaterals in magenta). Each point represents a single axon in one fish (control n= 43, DomA n=45). Error bars represent standard error of the mean. Statistical significance was determined using a one-way analysis of variance (ANOVA) *** = p < 2.2e-16. Scale bars = 100 μm.

By using the transgenic line *Tg*(*cntn1b:EGFP-CAAX*), we were then able to examine the axon collaterals that branch from the main caudal primary motor neuron axon. While staining with α-acetylated tubulin showed no differences in the presence of the main branches in primary motor neurons (Fig. 8D), tracings of the motor neuron axon collaterals in DomA-treated *Tg*(*cntn1b:EGFP-CAAX*) larvae showed a significant reduction in the length of the branches (Fig. 8E, 8F) (Control (mean ± SE) = 1457 μm ± 43, DomA (mean ± SE) = 527 μm ± 29, F(1, 74.6)= 315.87, p <2.2 e-16).

## DISCUSSION

Developmental exposure to DomA leads to persistent behavioral deficits, but the cellular and molecular mechanisms that underlie these effects are largely unknown. We used the zebrafish model and its well-characterized startle response circuit as tools to identify the cell types that are preferentially targeted by developmental DomA exposure and link these to sensorimotor processing deficits that occur during startle. Our results reveal that DomA exposure alters sensorimotor function, reducing responsiveness to stimuli and disrupting kinematics during startle. By imaging the neuronal components of the circuit, we found that DomA exposure led to the loss of reticulospinal neurons and to the reduction in axon collaterals in caudal primary motor neurons, while not affecting the presence of sensory ganglia or other hindbrain axon tracks. Taken together, these results suggest that DomA exposure leads to the loss of specific cells that are necessary for startle circuit function, resulting in behavioral startle deficits.

### Mauthner cells are targeted by DomA exposure

DomA prevents specific types of startle response from occurring in a manner consistent with the loss of a type of hindbrain reticulospinal neuron known as the Mauthner cell. The Mauthner cell is required for performing both electric field-induced startle and A/V-induced SLC startles (but not LLC startles) (Kohashi and Oda, 2008; Marsden and Granato, 2015; Tabor et al., 2014). Behaviors that require the Mauthner cell-mediated startle were lost in a majority of the DomA-exposed larvae. Furthermore, DomA exposed larvae preferentially performed LLC startles versus SLC startles even with high intensity A/V stimuli. These behavioral results suggest that the Mauthner cell is disrupted by DomA. Imaging results supported this; using two complementary approaches (spinal backfills and antibody staining for the neurofilament 3A10) we confirmed that the Mauthner cells were absent in a majority of DomA-treated larvae.

### DomA-induces motor axon collateral loss

DomA-exposed larvae showed kinematic deficits such as smaller bend angles and slower Mavs when performing all forms of startle tested (SLC, LLC, and electric field-induced startles). We hypothesized that this shared phenotype (altered startle kinematics) may be a result of disruptions to a cell type that is common in the neural circuits of these three types of startle responses. One of the shared neuronal components is the downstream primary motor neuron that activates the trunk and tail muscles required for the startle responses. While the main caudal primary motor neuron axons were present in DomA-exposed fish, the collaterals of these motor neuron axons were severely reduced. The reduction in the collaterals translated to fewer synapses to the trunk muscles, would could consequently have led to the observed shallower bend angles (McLean and Dougherty, 2015).

### Sensory system is an unlikely primary target for DomA

Developmental DomA exposure does not completely disrupt sensory processing during startle, consistent with the morphological data showing intact sensory ganglia. The sensory system is bypassed following direct-electric field stimulation, and yet DomA-exposed larvae still exhibited kinematic deficits under these conditions. This result indicates that the observed behavioral deficits cannot be due to solely deficits in the sensory system. Furthermore, while DomA-exposed larvae responded less often than control larvae, they still had the capacity to respond to A/V stimuli. Their responsiveness increased with increasing stimulus intensities, suggesting that DomA-exposed larvae have the ability to encode graded sensory information that translates to different degrees of responsiveness. Indeed, when assessing different components of sensory circuit, we found that DomA did not reduce the number of neuromasts or disrupt the presence of important sensory ganglia necessary for perceiving A/V stimuli, supporting the idea that DomA may not disrupt sensory system structures.

Importantly, we only assessed the presence versus absence of the specific sensory structures rather than their function. Previous studies have shown that the application of the ionotropic glutamate receptor agonist, AMPA, reduced the firing of afferent sensory neurons and the neurons’ responsiveness to hair cell activation (Sebe et al., 2017). Furthermore, we only assessed gross morphological features: we quantified the number of neuromasts present rather than more detailed structural features such as number of hair cells within a neuromast or the number of branches in afferent neuron terminals that synapse to the hair cells. Indeed, zebrafish hair cells express AMPA and KA receptors to which DomA binds (Hoppmann et al., 2008), making them potential targets for toxicity. Exposure to high doses of ionotropic glutamate agonists have been shown to lead to both the loss of hair cells within a single neuromast (with 300 μM KA or AMPA) (Sheets, 2017), and to the swelling of hair cell afferent synapses (with 100 μM AMPA) (Sebe et al., 2017).

It is conceivable that DomA may bind to ionotropic glutamate receptors in hair cells and at the mature hair cell ribbon synapse. DomA-induced overactivation of these receptors could then lead to hair cell death with individual neurons, and to the reduced firing capacity of afferent sensory neurons. Both of these events could contribute to the reduced responsiveness observed in DomA-exposed larvae. However, a more detailed analysis of individual hair cells and hair cell afferent synapse following DomA exposure would be necessary to determine whether this occurs.

Taken together, these results indicate that DomA does not alter the overall structures in the sensory system, nor does it affect the ability of larvae to increase their responsiveness with higher stimulus intensities. However, DomA-exposed fish are generally less responsive relative to control fish, which may be due to subtle perturbations to sensory systems that have yet to be investigated.

### Behavioral deficits are correlated with the severity of myelin phenotypes

The startle circuit is heavily myelinated to achieve rapid signaling. We showed previously that myelination deficits result from developmental exposure to DomA (Panlilio et al., 2020). By imaging *Tg*(*mbp:EGFP-CAAX*) larvae prior to behavioral tests, we were able to correlate the degree of myelin severity to the behavioral phenotypes.

We found that fish with more severe axonal and myelin defects were also likely to have more severe startle deficits. EGFP expression in *Tg*(*mbp:EGFP-CAAX*) fish was a proxy for the degree (score) of axonal and myelin defects because its expression is dependent both on intact myelin and on the presence of the axons (with no axons, there is nothing for oligodendrocytes to myelinate). Imaging results (Fig. 6 and 7) indicated that by the larval stages, reticulospinal neurons are lost in DomA-exposed fish, suggesting that the aberrant EGFP expression at these later developmental stages is due to the loss of axons. However, we cannot exclude the possibility that the loss of these axons was secondary to, or exacerbated by, the myelination defects. Our previous work (Panlilio et al., 2020) showed that as early as 2.5 dpf - the time at which axons are first wrapped – myelination is already perturbed. Thus, future studies are necessary to see whether DomA targets the axons first, followed by the myelination, or whether the loss of myelin leads to the axonal defects later.

Using EGFP labeling of myelin as a proxy for axonal and myelin integrity, we were able to determine that behavioral deficits observed were largely correlated to the severity in axon and myelin defects. Indeed, DomA-exposed larvae with “control-like” axons and myelin sheaths had fewer kinematic deficits relative to DomA-exposed fish with axon or myelin defects. However, there were some behavioral attributes for which this was not true. Fish with “control-like” axons and myelin did not always have “control-like” kinematics, especially during SLC startles. There are two potential explanations for this. 1) Fish that had “control-like” myelin sheaths and axons may have subtle myelin sheath and axonal defects that we could not distinguish from controls at the resolution we were imaging. These subtle defects could contribute to some of the observed SLC kinematic deficits. 2) Alternatively, DomA-exposed fish had myelin sheaths and axons that were indistinguishable from controls, but may have had other neuronal defects that we did not assess in relation to the behavioral changes. To test the latter scenario, future work could be done to assess the subset of the fish that had “control-like” myelin sheaths for effects on other cell targets, including motor axon collateral branching and ionotropic glutamate receptor activity in hair cells.

### DomA is not just a general neurotoxin

More generally, these findings suggest that DomA isn’t simply a nonspecific neurotoxin that targets all neural precursors and neurons, but instead one that targets specific neuronal and glial subtypes. The most apparent targets of DomA were the reticulospinal neurons in the hindbrain (including the Mauthner cell) and the caudal primary motor neurons. DomA had no apparent effects on other neurons and neuronal precursors examined including the medial longitudinal fasciculus, the hindbrain axonal tracts, the Rohon-beard sensory cell bodies, or the neural precursors in the spinal cord. While both Mauthner cells and caudal primary motor neurons express the ionotrophic glutamate receptors to which DomA binds (Patten and Ali, 2007; Todd et al., 2004; Warp et al., 2012), so too do other neurons and glial cells (Hoppmann et al., 2008), suggesting that other factors may be important in mediating susceptibility of subclasses of neurons.

The sensitivity of hindbrain reticulospinal neurons and motor neurons to DomA may be due to their intrinsic properties. Both of these neuron types have large axons, some with extensive axonal arbors that are located in either the spinal cord or periphery (Kimmel et al., 1982; Myers, 1985). Any perturbation that disrupts homeostatic mechanisms, such as slowing of axonal transport, could have important effects on these neurons, making these cells more susceptible to disease and environmental insults. In fact, motor neurons are known for being selectively vulnerable to insult because they are large, continuously active and heavily rely on mitochondrial respiration processes (Lewinski and Keller, 2005).

### Implications for human health

Finally, while these studies were done in zebrafish, these findings have implications for human health. Behavioral data indicate that DomA exposure in fish disrupts motor control. Motor deficits have been previously characterized from both incidental human exposures and animal exposure models (Shiotani et al., 2017; Teitelbaum et al., 1990; Wang et al., 2000). Adult humans acutely exposed to DomA developed sensorimotor neuropathy and axonopathy as assessed by electromyography (Teitelbaum et al., 1990). A subset of primates exposed orally at or near the accepted daily tolerable dose of 0.075 mg/kg developed visible hand tremors (Petroff et al., 2019). Rodents prenatally exposed to DomA (PND 10-17) developed aberrant gait patterns (Shiotani et al., 2017). These results suggest that motor deficits may be an important functional endpoint for DomA toxicity; future studies should assess both reflex and fine motor skills. One recent study in non-human primates found that *in utero* exposure to DomA had no impact on early survival reflexes in neonates (Grant et al., 2019). It would be useful to continue to trace motor skill development in these neonates to determine whether there are any potential latent effects, or whether more subtle motor skill deficits emerge.

The study also provides insights into potential cell targets for developmental exposures to DomA, demonstrating that DomA selectively targets specific neurons – the primary motor neurons and reticulospinal neuron subclasses. While humans do not have Mauthner cells, the intrinsic characteristics that make this cell type more vulnerable to toxins may be shared with other neurons in humans. Identifying these general characteristics (large axons, extensive arbors, location in the spinal cord) may also provide some useful hints as to which neurons are targeted in humans. While the identification of other candidates would require other animal models, our results using the zebrafish startle response can help guide further investigation of the cellular and molecular mechanisms underlying the behavioral deficits caused by early life exposure to DomA.

## CONCLUSION

This study characterized the effects of developmental DomA exposure on the startle response. The startle response circuit is well characterized, and the neural circuits that drive it are well known. Utilizing this knowledge, we were able to identify specific neural populations that may be more sensitive to early life exposure to DomA. Furthermore, this study illustrates the potential of using the startle response circuit as a tool to identify neuronal populations targeted by toxin or toxicant exposures.

## ACKNOWLEDGEMENTS

We thank H.E. Rivera and A.R. Solow for their advice on the statistical analysis, H.A. Burgess for providing the Flote software for startle kinematic analysis, B.G. Merrick for his help designing and building the startle apparatus, and N.R. Brun and L. Kerr for microscopy training and advice (MBL microscopy facility). We also thank the labs who generously provided us with zebrafish transgenic lines to make this work possible: B. Appel (University of Colorado, Denver), K.R. Monk (Oregon Health & Science University), C.K. Kaufman (Washington University School of Medicine in St. Louis), L. Zon (Harvard University), S. Kucenas (University of Virginia), and D.A. Lyons (University of Edinburg).

This research was supported by a WHOI Von Damm and Ocean Ridge Initiative Fellowships to J.M.P. and the Woods Hole Center for Oceans and Human Health (NIH: P01ES021923 and P01ES028938; NSF: OCE-1314642 and OCE-1840381). This paper includes data submitted by J.M.P. in partial fulfillment of the requirements for the degree of Doctor of Philosophy in Oceanography and Applied Ocean Science and Engineering at the Massachusetts Institute of Technology and the Woods Hole Oceanographic Institution (Panlilio, 2019)

