## Supplemental materials for "Developmental exposure to domoic acid disrupts startle response behavior and circuitry"

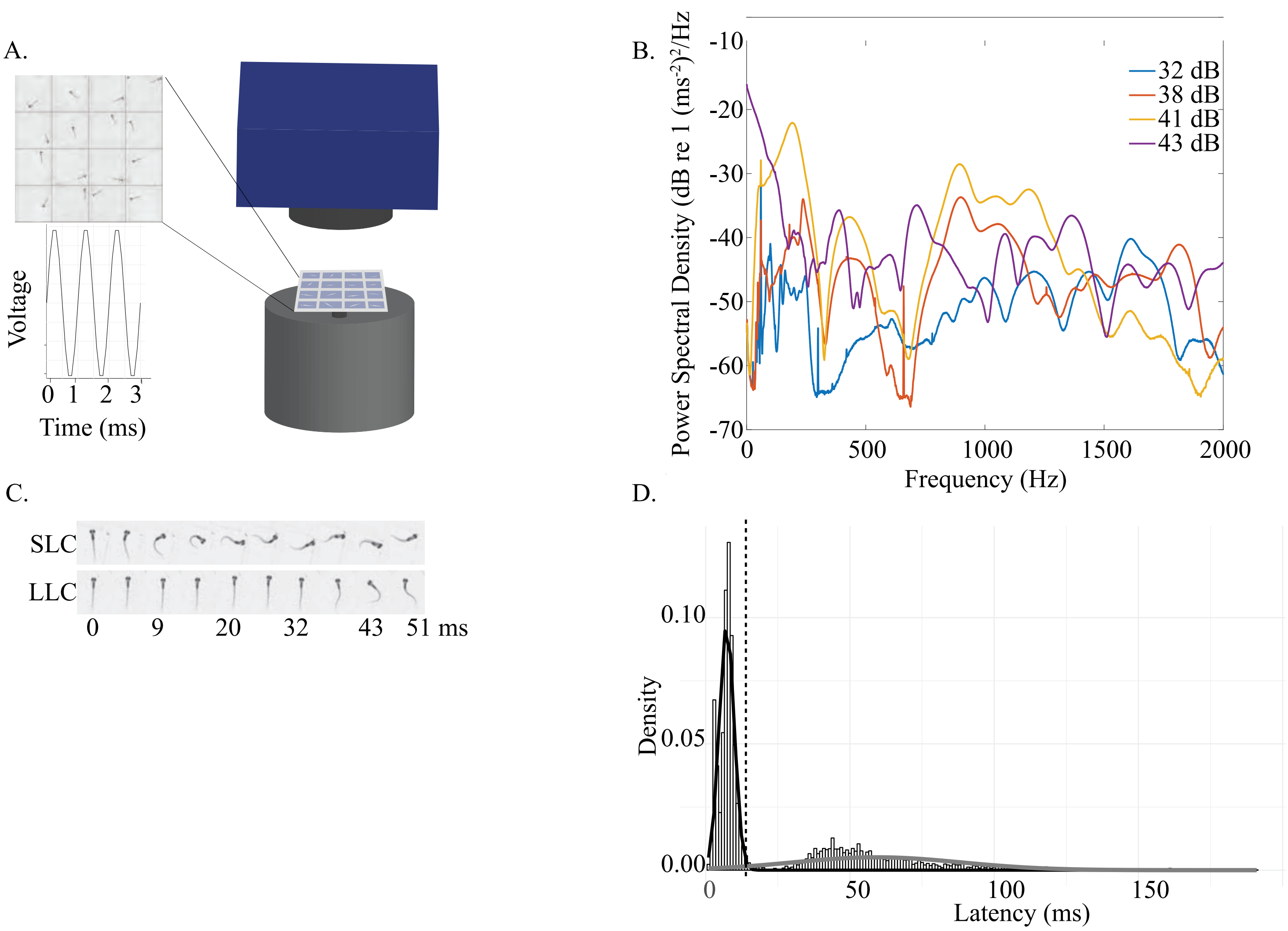


**Supplemental Fig 1: Startle behavioral apparatus and behavioral classification**

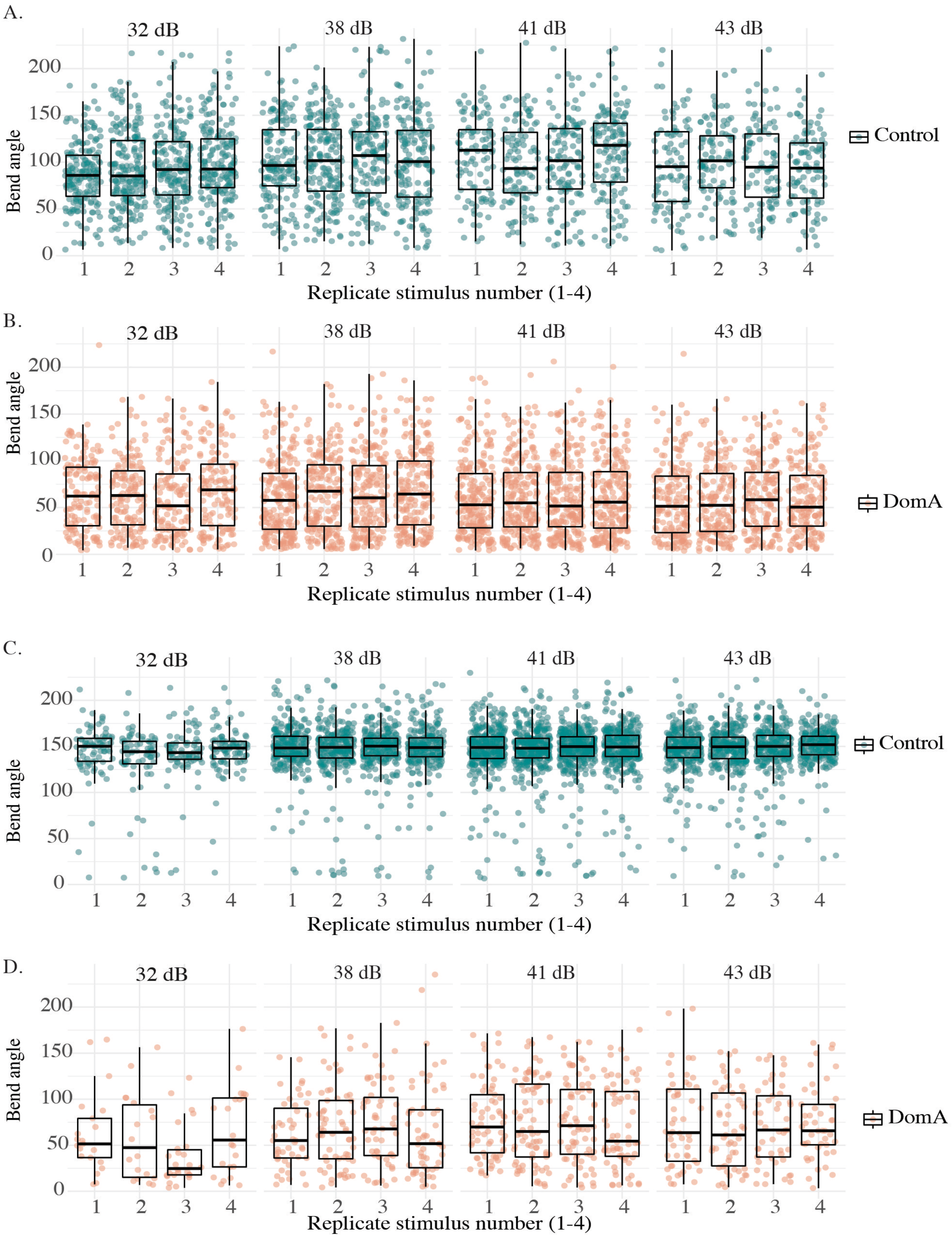


**Supplemental Fig 2: SLC and LLC startle bend angles from replicate stimuli**

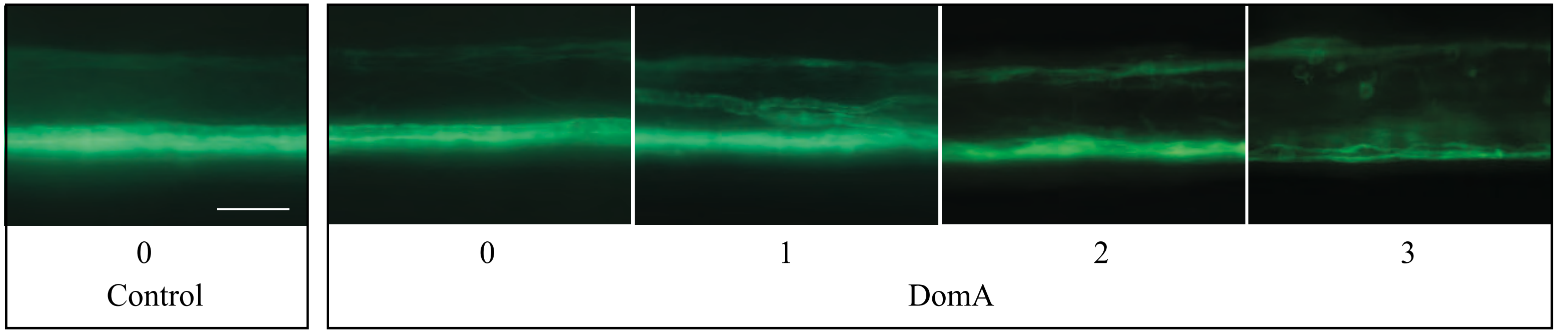


**Supplemental Figure 3: Range of classified myelin phenotypes.**
Myelin phenotypes were divided into three categories (0-3). 0 = control-like myelin sheaths to 3 = most severe phenotypes seen.

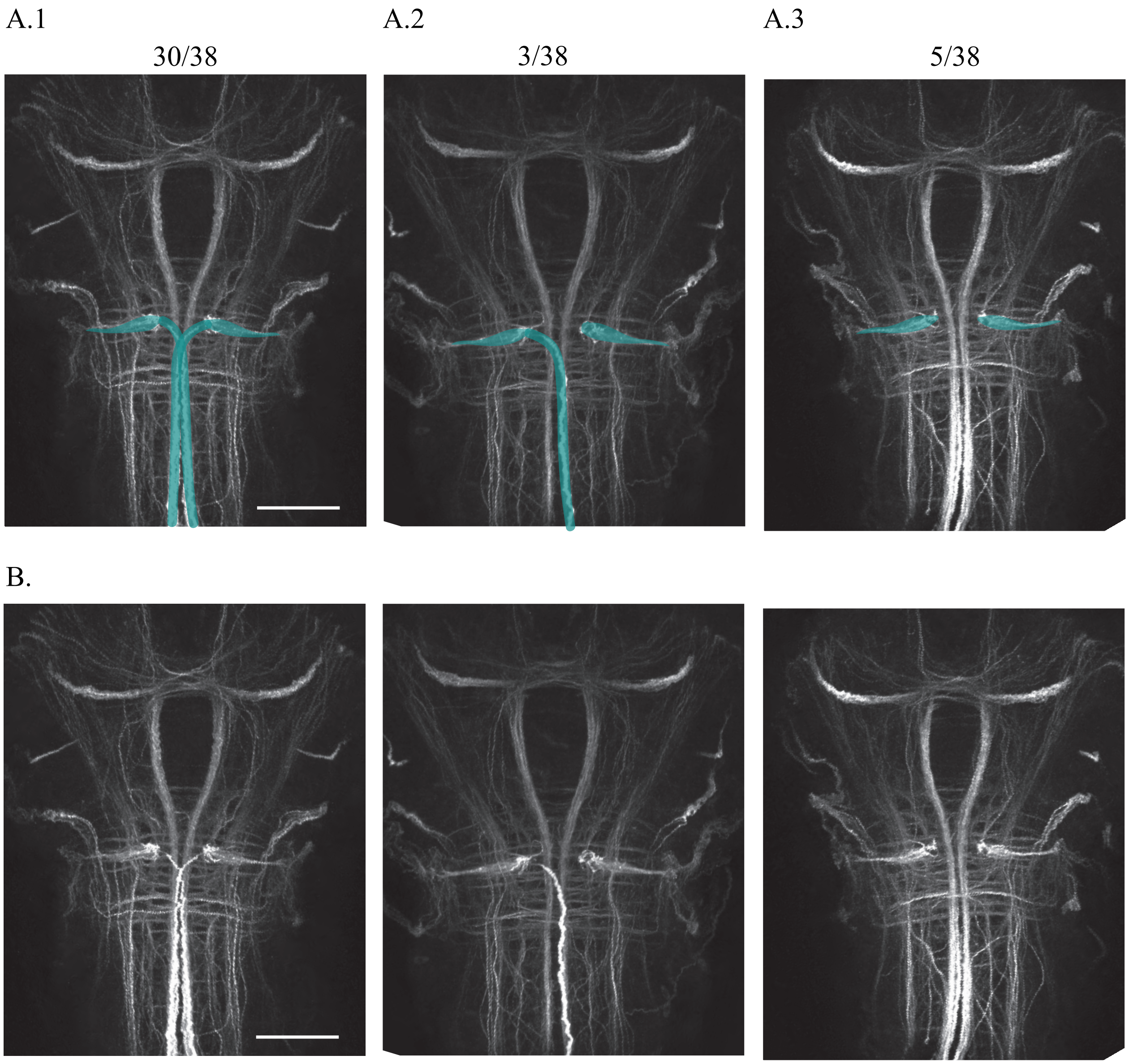


**Supplemental Figure 4: Range of 3A10 staining observed in control larvae**
**(A)** Larval brains were dissected then stained using the 3A10 antibody. In a subset of the control animals, the Mauthner cell bodies were present, but one (A2) or both (A3 and A4) of the Mauthner axons were absent, likely as an artifact of the dissections. Mauthner cell bodies and axons are highlighed in teal. Ratios above each image show the number of control fish with each of these phenotypes. Scale bar = 100 μm.
**(B)** The same images without the highlighted regions.

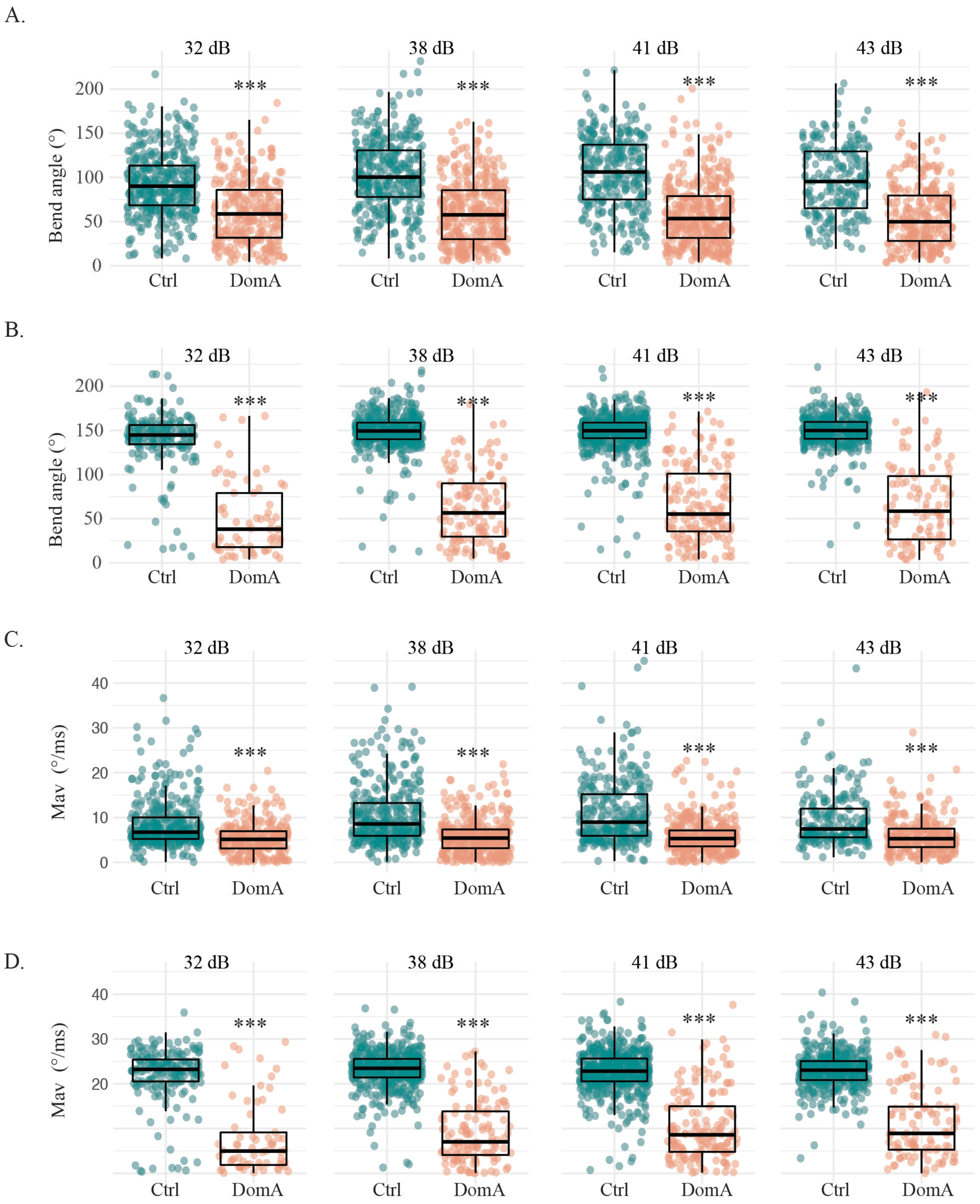


**Supplemental Fig 5: Domoic acid-exposed larvae have reduced bend angles and maximal angular velocities across all stimulus intensities tested when given auditory/vibrational stimuli.**

**SUPPLEMENTAL INFORMATION**

**Supplemental Fig 1: Startle behavioral apparatus and behavioral classification**

**(A)** Diagram of apparatus used to assess startle responses to A/V stimuli. A vibrational exciter delivers a 3 millisecond, 1000 Hz pulse to 16-well plate that contains single larvae within individual wells. The amplitude of the pulse was varied to deliver a range of stimulus intensities. A high-speed camera captured startle responses at 1000 frames per second.

**(B)** Power spectrum of the four Auditory/vibrational stimuli intensities provided (32, 38, 41, and 43 dB).

**(C)** Temporal projections of the Mauthner-dependent short latency c-startle response (SLC) which initiates within 14ms or less after the delivery of the stimuli. Temporal projections of the long latency c-startle response (LLC) which initiates more than 14 milliseconds after the stimulus.

**(D)** Density histogram of the latency distribution for control fish. Overlaid are two Gaussian curves that were fit for the data. The black curve represents SLC responses, and grey represents LLC response. Mixture. The dashed black vertical line at 14 milliseconds represents the cut-off by which there is a greater than 50% probability of a given data point belonging to either the modeled SLC or LLC distribution.

**Supplemental Fig 2: SLC and LLC startle bend angles from replicate stimuli**

(**A)** Bend angle during LLC-type startles for control fish across the replicate stimuli (1-4 within each intensity (32 dB, 38 dB, 41 dB, 43 dB)

**(B)** Bend angle during LLC-type startles for DomA-exposed fish across the replicate stimuli (1-4 within each intensity (32 dB, 38 dB, 41 dB, 43 dB)

**(C)** Bend angle during SLC-type startles for control fish across the replicate stimuli (1-4 within each intensity (32 dB, 38 dB, 41 dB, 43 dB)

**(D)** Bend angle during SLC-type startles for DomA-exposed fish across the replicate stimuli (1-4 within each intensity (32 dB, 38 dB, 41 dB, 43 dB)

**Supplemental Fig 3: Range of classified myelin phenotypes**

**(A)** Myelin phenotypes were divided into three categories (0-3). 0 = control-like myelin sheaths, 3 = most severe phenotypes seen.

**Supplemental Fig 4: Range of phenotypes observed in control larvae**

**(A)** Larval brains were dissected then stained using the 3A10 antibody. In a subset of the control animals, the Mauthner cell bodies were present but one (A2) or both (A3 and A4) of the Mauthner axons were absent, likely as an artifact of the dissections. Mauthner cell bodies and axons are highlighed in teal. Ratios above each image show the number of control fish with each of these phenotypes.

**(B)** The same images without the highlighted regions.

**Supplemental Fig 5: Domoic acid-exposed larvae have reduced bend angles and maximal angular velocities across all stimulus intensities tested when given auditory/vibrational stimuli.**

- **(A)** Bend angles during LLC-type startles with increasing stimulus intensities
- **(B)** Maximal angular velocities during LLC-type startles with increasing stimulus intensities
- **(C)** Bend angles during SLC-type startles with increasing stimulus intensities
- **(D)** Maximal angular velocities during SLC-type startles with increasing stimulus intensities

Individual points represent the median kinematic response of a single fish for up to 4 replicate stimuli. Asterisks represent statistical significance between DomA and controls determined using an Aligned Ranked Transformed ANOVA test (*** $p < 1e{-6}$).

**Supplemental Tables**

**Table S1: Stimulus regime for individual behavioral trials**

| trial number | stimulus regime in decibels (# of times stimulus is given) | dpf tested | Myelin assessment |
|---|---|---|---|
| 1 | 32 (x4), 38 (x4), 41 (x4) | 6 dpf | no |
| 2 | 32 (x4), 38 (x4), 41 (x7) | 6 dpf | no |
| 3 | 32 (x4), 38 (x4), 41 (x4), 43 (x4) | 6 dpf | yes |
| 4 | 32 (x4), 38 (x4), 41 (x4), 43 (x4) | 6 dpf | yes |
| 5 | 32 (x4), 38 (x4), 41 (x4), 43 (x4) | 6 dpf | yes |
| 6 | 32 (x4), 38 (x4), 41 (x4), 43 (x4) | 6 dpf | yes |
| 7 | 32 (x4), 38 (x4), 41 (x4), 43 (x4) | 6 dpf | yes |
| 8 | 32 (x4), 38 (x4), 41 (x4), 43 (x4) | 6 dpf | yes |
| 9 | 32 (x4), 38 (x4), 41 (x4), 43 (x4) | 6 dpf | yes |

**Table S2: Percentage of fish that performed LLCs, SLCs, or did not respond within a given treatment and to a selected stimulus intensity**

| | Control | | | | DomA | | | |
|---|---|---|---|---|---|---|---|---|
| Stimulus Intensity | % LLC | % NR | % SLC | N | % LLC | % NR | % SLC | N |
| 32 dB | 31.9% | 54.2% | 13.9% | 583 | 30.2% | 65.2% | 4.6% | 460 |
| 38 dB | 29.1% | 21.9% | 49.0% | 567 | 47.8% | 41.9% | 10.4% | 454 |
| 41 dB | 19.9% | 12.7% | 67.4% | 528 | 51.6% | 33.0% | 15.5% | 446 |
| 43 dB | 22.3% | 10.3% | 67.4% | 417 | 48.5% | 37.1% | 14.4% | 361 |

*Notes*: N= total number of fish tracked per each stimulus intensity combining all trials together. NR= no response, SLC= Short latency c-bend startles, LLC= Long latency c-bend startles

**Table S3: Generalized linear mixed-effects model for responsiveness in DomA and Control fish**

| decibel level | | Estimate | Std. Error | Z value | P value |
|---|---|---|---|---|---|
| **32** | (Intercept) | -0.068 | 0.168 | -0.404 | 0.6863 |
| | DomA | -0.474 | 0.191 | -2.479 | 0.0132 |
| **38** | (Intercept) | 1.387 | 0.222 | 6.246 | 4.20E-10 |
| | DomA | -0.902 | 0.194 | -4.640 | 3.49E-06 |
| **41** | (Intercept) | 2.252 | 0.237 | 9.492 | < 2e-16 |
| | DomA | -1.381 | 0.233 | -5.923 | 3.15E-09 |
| **43** | (Intercept) | 2.351 | 0.322 | 7.302 | 2.83E-13 |
| | DomA | -1.733 | 0.341 | -5.088 | 3.61E-07 |

*Note*: Intercept is control fish injected with saline.

**Table S4: Generalized linear mixed-effects model for performing SLC over LLC startles**

| decibel level | | Estimate | Std. Error | Z value | P value |
|---|---|---|---|---|---|
| **32** | (Intercept) | -0.975 | 0.1704 | -5.718 | 1.08E-08 |
| | DomA | -1.076 | 0.1379 | -7.807 | 5.84E-15 |
| **38** | (Intercept) | 0.349 | 0.187 | 1.865 | 6.21E-02 |
| | DomA | -1.960 | 0.304 | -6.455 | 1.08E-10 |
| **41** | (Intercept) | 0.999 | 0.172 | 5.819 | 5.91E-09 |
| | DomA | -2.425 | 0.408 | -5.942 | 2.82E-09 |
| **43** | (Intercept) | 0.963 | 0.212 | 4.539 | 5.65E-06 |
| | DomA | -2.500 | 0.512 | -4.878 | 1.07E-06 |

*Note*: Intercept is control fish injected with saline.

**Table S5: Generalized linear mixed-effects model for performing SLC over LLC startles in Control and DomA fish that are subcategorized by myelin phenotype (from a 43 dB A/V stimulus)**

| Treatment | Myelin Category | Estimate | Std. Error | Z value | P value |
|---|---|---|---|---|---|
| Intercept (Control) | 0 | 0.958 | 0.213 | 4.489 | 7.15E-06 |
| DomA | 0 | 0.225 | 0.548 | 0.411 | .0681 |
| | 1 | -1.552 | 0.442 | -3.51 | 0.0004 |
| | 2 | -1.633 | 0.485 | -3.366 | 0.0008 |
| | 3 | -3.196 | 0.424 | -7.537 | 4.81E-14 |

*Notes*: Intercept is control fish injected with saline. Numbers 0-3 denote myelin categories into which fish were classified prior to behavioral analyses. 0 = phenotype indistinguishable from Control to 3= most severe phenotype observed. See *Methods* for details

**Table S6: Nonparametric multiple comparison tests for SLC startle kinematics by stimulus intensity**

| | | | Bend angle | | Mav | |
|---|---|---|---|---|---|---|
| **Stimulus intensity** | **N Treatment (category)** | **Comparison** | **Est [Lower, Upper]** | **p** | **Est [Lower, Upper]** | **p** |
| 32 dB | Control(0) = 153, DomA(0) = 5, DomA(1) = 7, DomA(2) = 7, DomA(3) = 28 | Control (0), DomA (0) | 0.058 [0.006, 0.109] | < 1 e -16 | 0.251 [-0.451, 0.9523 | 5.44 e-01 |
| | | Control (0), DomA (1) | 0.072 [0.007, 0.137] | < 1 e -16 | 0.225 [-0.244, 0.694] | 2.22 e-01 |
| | | Control (0), DomA (2) | 0.041 [-0.004, 0.086] | < 1 e -16 | 0.052 [-0.019, 0.123] | 2.24 e-7 |
| | | Control (0), DomA (3) | 0.023 [-0.002, 0.048] | < 1 e -16 | 0.044 [-0.014, 0.101] | 1.98e-9 |
| 38 dB | Control(0) = 416, DomA(0) = 16 , DomA(1) = 22, DomA(2) = 8, DomA(3) = 47 | Control (0), DomA (0) | 0.094 [-0.037, 0.225] | 3.45e -07 | 0.187 [-0.004, 0.377] | 0.0013 |
| | | Control (0), DomA (1) | 0.036 [0.001, 0.070] | < 1 e -16 | 0.052 [0.007, 0.097] | < 1 e -16 |
| | | Control (0), DomA (2) | 0.011 [-0.003, 0.025] | < 1 e -16 | 0.006 [-0.004, 0.016] | < 1 e -16 |
| | | Control (0), DomA (3) | 0.004 [-0.002, 0.010] | < 1 e -16 | 0.009 [-0.004, 0.021] | < 1 e -16 |
| 41 dB | Control(0) = 492, DomA(0) = 20, DomA(1) = 35, DomA(2) = 9, DomA(3) = 51 | Control (0), DomA (0) | 0.174 [0.009, 0.339] | 0.0002 | 0.311 [0.096, 0.526] | 9.80 e-02 |
| | | Control (0), DomA (1) | 0.034 [0.006, 0.062] | < 1 e -16 | 0.168 [0.032, 0.303] | 7.70 e-06 |
| | | Control (0), DomA (2) | 0.017 [0.000, 0.034] | < 1 e -16 | 0.015 [0.00, 0.029] | < 1 e -16 |
| | | Control (0), DomA (3) | 0.015 [0.001, 0.029] | < 1 e -16 | 0.028 [-0.01, 0.067] | < 1 e -16 |
| 43 dB | Control(0) = 425, DomA(0) = 18, DomA(1) = 34, DomA(2) = 12, DomA(3) = 37 | Control (0), DomA (0) | 0.153 [-0.010, 0.316] | 1.53 e -04 | 0.372 [0.097, 0.647] | 5.87 e-01 |
| | | Control (0), DomA (1) | 0.026 [-0.004, 0.056] | < 1 e -16 | 0.104 [-0.014, 0.195] | 5.08 e -12 |
| | | Control (0), DomA (2) | 0.094 [-0.147, 0.334] | < 1 e -16 | 0.018 [-0.001, 0.046] | < 1 e -16 |
| | | Control (0), DomA (3) | 0.023 [-0.036, 0.082] | < 1 e -16 | 0.032 [-0.032, 0.097] | < 1 e -16 |

*Notes*: Category = myelin category number, Est = estimator, [Lower, Upper] = Lower and upper limits of the 95% confidence intervals. Significant values are colored in red.

**Table S7: Nonparametric multiple comparison tests for LLC startle kinematics by stimulus intensity**

| Stimulus intensity | N | Comparison | Bend angle: Estimator [Lower, Upper] | Bend angle: p.Value | Mav: Estimator | Mav: p.Value |
|---|---|---|---|---|---|---|
| 32 dB | Control(0) = 353, DomA(0) = 10, DomA(1) = 28, DomA(2) = 15, DomA(3) = 158 | Control (0), DomA (0) | 0.390 [0.083, 0.698] | 7.31 e-1 | 0.418 [0.127, 0.71] | 8.59 e -1 |
| | | Control (0), DomA (1) | 0.418 [0.222, 0.614] | 6.17 e-1 | 0.505 [0.315, 0.695] | 9.99 e-1 |
| | | Control (0), DomA (2) | 0.344 [0.092, 0.597] | 2.95 e-1 | 0.373 [0.156, 0.591] | 3.39 e-1 |
| | | Control (0), DomA (3) | 0.242 [0.170, 0.314] | 1. 74 e-6 | 0.271 [0.197, 0.346] | 2.60 e-5 |
| 38 dB | Control(0) = 329, DomA(0) = 10 , DomA(1) = 39, DomA(2) = 19, DomA(3) = 240 | Control (0), DomA (0) | 0.361 [0.160, 0.562] | 2.23 e-1 | 0.426 [0.205, 0.647] | 7.77 e-1 |
| | | Control (0), DomA (1) | 0.308 [0.178, 0.437] | 4.51 e-3 | 0.381 [0.252, 0.510] | 7.26 e-2 |
| | | Control (0), DomA (2) | 0.246 [0.088, 0.403] | 2.39 e-3 | 0.250 [0.100, 0.401] | 2.20 e-3 |
| | | Control (0), DomA (3) | 0.190 [0.137, 0.243] | 9.88 e-15 | 0.232 [0.174, 0.291] | 9.85 e-9 |
| 41 dB | Control(0) = 276 DomA(0) = 10, DomA(1) = 41, DomA(2) = 18, DomA(3) = 253 | Control (0), DomA (0) | 0.352 [0.011, 0.693] | 5.81 e-1 | 0.434 [0.123, 0.744] | 9.36 e-01 |
| | | Control (0), DomA (1) | 0.286 [0.161, 0.411] | 2.25 e-3 | 0.361 [0.236, 0.486] | 2.97 e-2 |
| | | Control (0), DomA (2) | 0.207 [0.047, 0.367] | 1.41 e-3 | 0.243 [0.094, 0.392] | 1.99 e-3 |
| | | Control (0), DomA (3) | 0.154 [0.104, 0.205] | 2.96 e-12 | 0.202 [0.144, 0.259] | 4.07 e-10 |
| 43 dB | Control(0) = 219, DomA(0) = 7, DomA(1) = 39, DomA(2) = 20, DomA(3) = 219 | Control (0), DomA (0) | 0.297 [-0.087, 0.682] | 3.60 e-1 | 0.379 [0.067, 0.691 | 0.629 |
| | | Control (0), DomA (1) | 0.326 [0.172, 0.480] | 3.01 e-2 | 0.397 [0.238, 0.557] | 0.239 |
| | | Control (0), DomA (2) | 0.283 [0.114, 0.452] | 1.71 e-2 | 0.485 [0.241, 0.73] | 0.999 |
| | | Control (0), DomA (3) | 0.196 [0.127, 0.265] | 3.71 e-7 | 0.248 [0.174, 0.321] | 0.0003 |

*Notes*: Category = myelin category number, Est = estimator, [Lower, Upper] = Lower and upper limits of the 95% confidence intervals. Significant values are colored in red.

**Table S8: Nonparametric multiple comparison tests for startle kinematics following direct electric field stimulation**

| | | Bend angle | | Mav | |
|---|---|---|---|---|---|
| **N** | **Comparison** | **Estimator [Lower, Upper]** | **p.Value** | **Estimator** | **p.Value** |
| Control(0) = 74, DomA(0) = 6, DomA(2) = 5, DomA(3) = 14 | Control (0), DomA (0) | 0.39 [-0.15, 0.94] | 8.4 e -01 | 0.42 [-0.11, 0.95] | 9.2 e -1 |
| | Control (0), DomA (2) | 0.22 [-0.41, 0.86] | 3.90 e-01 | 0.16 [-0.25, 0.57] | 8.8 e -2 |
| | Control (0), DomA (3) | 0.05 [-0.11, 0.21] | 1.00 e-03 | 0.036 [-0.04, 0.11] | 1.3 e -7 |

*Notes*: Category = myelin category number, Est = estimator, [Lower, Upper] = Lower and upper limits of the 95% confidence intervals. Significant values are colored in red.
